# Brainwide hemodynamics predict EEG neural rhythms across sleep and wakefulness in humans

**DOI:** 10.1101/2024.01.29.577429

**Authors:** Leandro P. L. Jacob, Sydney M. Bailes, Stephanie D. Williams, Carsen Stringer, Laura D. Lewis

## Abstract

The brain exhibits rich oscillatory dynamics that play critical roles in vigilance and cognition, such as the neural rhythms that define sleep. These rhythms continuously fluctuate, signaling major changes in vigilance, but the widespread brain dynamics underlying these oscillations are difficult to investigate. Using simultaneous EEG and fast fMRI in humans who fell asleep inside the scanner, we developed a machine learning approach to investigate which fMRI regions and networks predict fluctuations in neural rhythms. We demonstrated that the rise and fall of alpha (8-12 Hz) and delta (1-4 Hz) power— two canonical EEG bands critically involved with cognition and vigilance—can be predicted from fMRI data in subjects that were not present in the training set. This approach also identified predictive information in individual brain regions across the cortex and subcortex. Finally, we developed an approach to identify shared and unique predictive information, and found that information about alpha rhythms was highly separable in two networks linked to arousal and visual systems. Conversely, delta rhythms were diffusely represented on a large spatial scale primarily across the cortex. These results demonstrate that EEG rhythms can be predicted from fMRI data, identify large-scale network patterns that underlie alpha and delta rhythms, and establish a novel framework for investigating multimodal brain dynamics.

**Author summary:** Neurons often fire in synchrony, generating rhythms that play major roles in brain functioning. These rhythms are hallmarks of different brain states of vigilance, such as sleep and wakefulness. Sleep disorders are extremely prevalent among adults, and studying neural rhythms associated with vigilance states is a key step towards understanding sleep disorders and how healthy sleep can be restored. Measuring how neural rhythms affect the brain, however, is difficult: the primary method used in humans, electroencephalography (EEG), can only measure neural activity close to the scalp. EEG can be combined with functional magnetic resonance imaging (fMRI), which is capable of measuring activity in deep brain regions, but fMRI data can be difficult to analyze, as it estimates neural activity indirectly by measuring changes in blood oxygenation. We developed an approach to analyze combined EEG-fMRI data using machine learning, and used it to investigate how fluctuations in neural rhythms across sleep and wakefulness are tied to changes in neural activity throughout the whole brain. Our results describe how different brain networks are coupled to alpha and delta rhythms, and provide a new approach for analyzing EEG-fMRI data that can be employed to investigate other neural rhythms necessary for healthy brain functioning.

## INTRODUCTION

As the brain navigates tasks and states, neurons frequently fire in synchrony, generating rich oscillatory dynamics with varied temporal and spatial distributions. These neural rhythms have been studied with scalp EEG for almost a century, and have formed a core aspect of modern neuroscience due to their established links to cognitive computations and brain states. Neural rhythms are particularly fundamental to the study of vigilance states and sleep, which are defined by the appearance of distinct oscillatory EEG patterns with well-studied cognitive outcomes ^1^, including several forms of memory ^2–10^ and attention ^11,12^, along with basic physiological processes such as brain waste clearance ^13^. Extensive work has been dedicated to identifying the oscillatory generators behind these electrophysiological rhythms. However, the waxing and waning of neural rhythms can reflect widespread variations in brain activity, beyond the specific subset of neurons oscillating at a given frequency. In this work, we developed a novel predictive approach that investigates the brainwide variations underlying neural rhythms using simultaneous EEG and fast fMRI. We focused on two widely-studied neural rhythms with key relevance for the study of arousal and cognition: delta (1-4 Hz) and alpha (8-12 Hz) oscillations. Alpha (Fig. 1a) is implicated in an array of cognitive processes ^12,14,15^, and maximally detected in occipital electrodes during eyes-closed wakeful rest. Delta (Fig. 1a) is most prominent during non-rapid eye movement (NREM) sleep and strongly linked to its cognitive benefits ^4,5,9,16^.

**Fig. 1:**
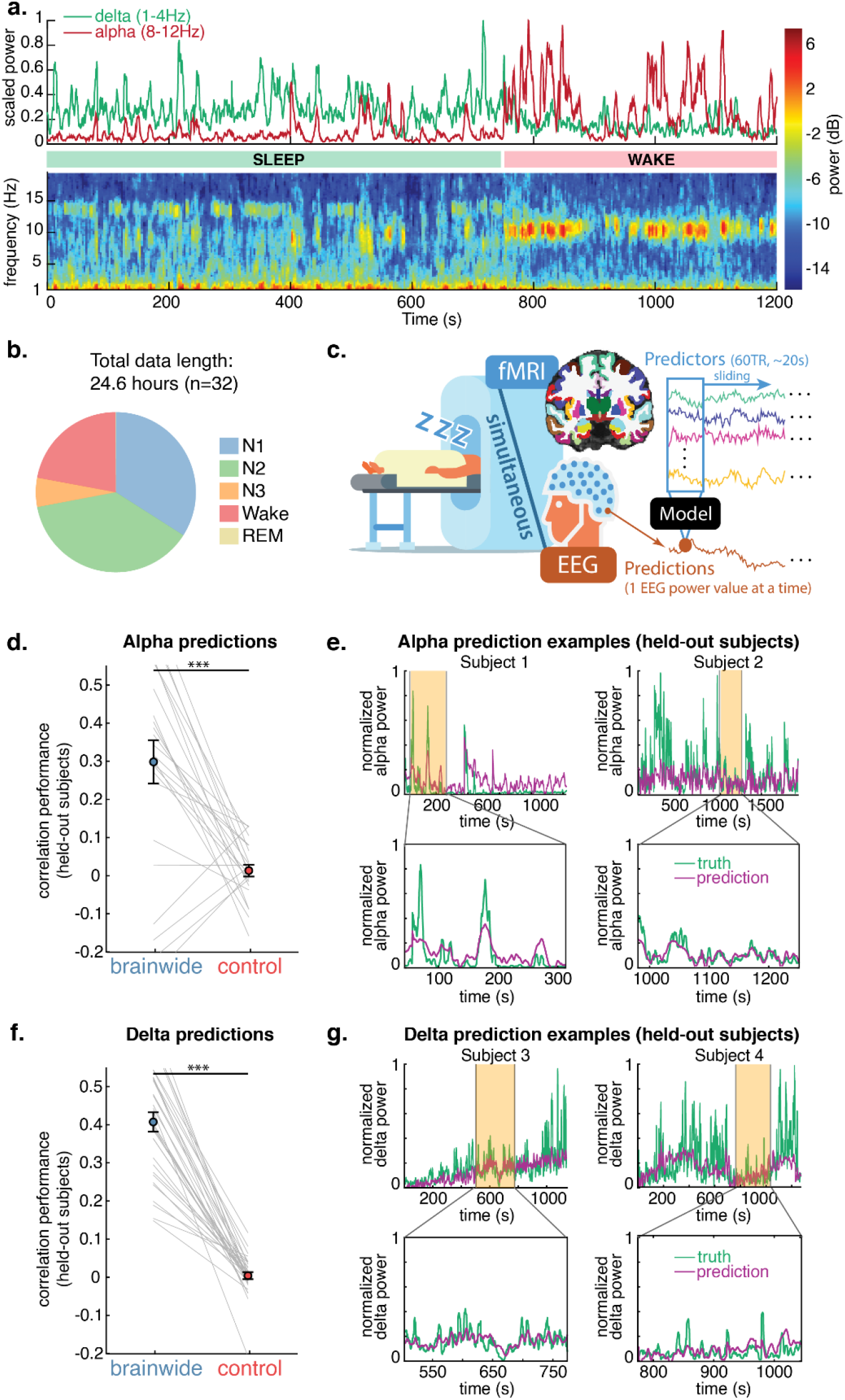
Machine learning can predict dynamic fluctuations in neural rhythms using brainwide fast fMRI timeseries. **a.** Example of occipital alpha and delta EEG power fluctuating in one subject. **b.** Sleep stage distribution shows subjects drifted between wake and sleep (primarily N1 and N2). **c.** EEG alpha and delta power were separately predicted from simultaneous fMRI data, collected while subjects rested inside the scanner with their eyes closed. fMRI data was parcellated into 84 cortical, subcortical, and non-gray matter regions. Each point in the EEG power series, interpolated to match TR times, was predicted by 60 TRs of parcellated fMRI data. **d.** Correlation between alpha predictions (on held-out subjects) and ground truth, demonstrating that predictions using brainwide fMRI data are significantly better than the control condition, in which the fMRI data was shuffled within each subject run. *** p<0.001, two-tailed paired t-test. Gray lines indicate held-out subject prediction performance. Error bars are SEM; N=24. **e.** Examples of alpha prediction in individual subjects, showing tracking of short- and long-timescale alpha power fluctuations. **f.** Delta prediction performance, across the same conditions as panel d, shows significantly above-chance delta prediction. N=32. **g.** Examples of delta prediction in individual subjects.

While electrophysiological studies have revealed extensive links between these rhythms and the cognitive processes they support, identifying brainwide dynamics coupled to neural rhythms is technically challenging: scalp EEG suffers from low spatial resolution and is unable to resolve deep brain activity, while invasive methods such as electrocorticography have limited spatial coverage. To obtain high spatial resolution and brainwide coverage, EEG can be acquired simultaneously with functional magnetic resonance imaging (fMRI). Alpha rhythms in particular have been widely studied using EEG-fMRI, and prior studies have identified visual and thalamic regions that are correlated to alpha ^17–26^. fMRI activity has also been linked to continuous delta power fluctuations ^27–29^, and prior work has identified both widespread correlations between delta and several fMRI brain regions ^28^, as well as correlations with higher-frequency fMRI signals ^29^. These prior studies show that fMRI signals covary meaningfully with EEG neural rhythms, demonstrating the potential of multimodal imaging to identify brainwide correlates of electrophysiological rhythms. Intriguingly, the presence of these correlations suggests that it might be possible to decode information about the EEG solely using fMRI signals, despite their much lower temporal resolution—however, this possibility has not yet been tested.

We aimed to decode EEG rhythms from fMRI activity in order to investigate which brainwide network dynamics underlie the distinctive changes in EEG rhythms that signal fluctuations in vigilance state. In particular, we hypothesized that additional features of fMRI signals might contain important dynamics related to EEG rhythms that may not be captured by traditional models. The BOLD (blood-oxygenation-level-dependent) signal measured by fMRI is an indirect measure of brain activity, and their coupling varies across brain areas ^30^, with studies showing that the standard HRF does not account for some temporal properties of fMRI signals ^31–33^. Approaches that fit a more flexible HRF, on the other hand, can overfit the data if cross-validation is not used ^34^. Predictive machine learning methods are thus a powerful approach for fMRI ^35,36^: they can learn complementary information across multiple brain regions with flexible data-driven functions, and they inherently involve cross-validation to ensure results are generalizable. These approaches have identified relationships between neural activity and fMRI signals ^37^, classified brain states ^38,39^, and predicted a continuous vigilance index from fMRI ^40,41^. We thus sought to determine whether continuous variations in EEG rhythms can be predicted from fMRI, and if this decoding approach can allow for the identification of distinct network dynamics underlying EEG rhythms.

We designed a predictive framework to identify how brainwide dynamics are linked to distinct EEG rhythms, using linear machine learning to predict alpha and delta power from fMRI data as a proof-of-concept. We hypothesized that accelerated fMRI acquisition could provide sufficient information to effectively predict fluctuations in EEG using fMRI data. We used two independent datasets of simultaneous EEG and accelerated fMRI (TR<400 ms), exploiting the high temporal information of our fMRI data to model fluctuations in neural rhythms across wakefulness and sleep. Critically, our goal was not to localize focal oscillatory sources, but rather to identify the brainwide activity patterns that are coupled to variations in EEG rhythms, as scalp oscillations can reflect brain dynamics that need not oscillate at the same frequency. We first show that we can successfully predict EEG neural rhythms from hemodynamics in subjects not seen during training. We find that distinct and replicable fMRI patterns predict each rhythm, and that predictive information is present even at the single-region level. Finally, we develop an analysis to probe the joint fMRI information across brain regions, and use it to identify distinct network patterns underlying each neural rhythm. This approach identifies how fMRI information predictive of neural rhythms is spatially distributed across the brain, separating networks of regions with shared (vs. distinct) information about EEG dynamics. We found a large, diffuse network of primarily cortical regions that predicted variations in delta power, whereas alpha power was predicted by two separable networks primarily representing the distinct dynamics of the visual system and subcortical arousal control circuits.

## RESULTS

### EEG alpha and delta power were predicted from brainwide fMRI data in held-out subjects

We first investigated whether it is possible to predict the continuous fluctuations of neural rhythms from simultaneously acquired fast fMRI data. Prior work successfully predicted EEG-based vigilance metrics (a ratio between two frequency bands) from simultaneous fMRI data and achieved a mean Pearson’s linear correlation value of *r*=0.31 between predictions and targets when training on the same subjects ^41^. However, predicting distinct rhythms is a more challenging task, as fluctuations in individual frequency bands are noisier and more complex than global variations in vigilance metrics, which collapse multiple distinct EEG properties into a single simplified metric. Additionally, our goal of generalizing to out-of-sample subjects could make predicting individual frequency bands particularly difficult, as EEG power magnitudes exhibit strong individual differences. Thus, based on prior work and the variability of individual EEG, we expected a performance ceiling in the range of *r*=0.3 (Pearson’s linear correlation between held-out subject predictions and ground truth). We hypothesized that the high temporal resolution of our fast fMRI data would allow us to predict variations in EEG rhythms at or above this performance ceiling.

We tested our models on held-out subjects (not seen during training) to ensure overfitting did not inflate our performance estimates and to assess the generalizability of the model across individuals. We employed two separate datasets collected at different sites, to further confirm reproducibility (dataset A: *n*=−21, *TR*=0.378s; dataset B: *n=*11, *TR*=0.367s). For alpha analyses, spectrograms were visually inspected to identify whether subjects had clear eyes-closed alpha rhythms (see Fig. a); prior work has shown that not all individuals exhibit increases in the alpha band during wake with eyes closed ^42–45^, and given that these individuals likely possess distinct alpha rhythm dynamics, we removed them from alpha training and analyses (*n=*3 and *n=*5 were removed from dataset A and B respectively only for alpha analyses).

We trained machine learning models to separately predict EEG alpha and delta power, calculated in sliding 5-second windows from occipital electrodes (which are less susceptible to MR-induced noise), from simultaneously collected fMRI data, in subjects drifting in and out of sleep (Fig. 1b-c). The fMRI data was parcellated in 84 anatomical regions, consisting of 62 cortical regions (31 in each hemisphere), 14 subcortical regions (7 in each hemisphere), and 8 non-gray matter regions (white matter and ventricles). The functional voxels in each parcel were averaged to create a mean time series for each region. Models were trained on sliding windows (length of 60 TRs, ~20s) of these parcellated fMRI data, with each segment yielding a single-point prediction of normalized EEG power (at the temporal center of the window). We centered the window of fMRI predictors on the EEG point being predicted to allow the model to flexibly learn whichever fMRI temporal patterns were most relevant, rather than assuming a specific lag between EEG and fMRI. Models were thus trained on 5,040 predictors (84 regions with 60 time points per region). In this proof-of-concept, we opted for a linear regression model, which learns one beta weight value for each predictor using stochastic gradient descent with ridge regularization (see Methods for details). Due to distinct imagining parameters, training was carried out separately for each dataset, iteratively on all subjects but one, with testing then performed on the held-out subject; predictive performance on held-out subjects was combined for statistical analyses and visualizations. The performance was compared to a control in which models were trained on fMRI data that was temporally shuffled (preserving the EEG structure, but breaking the relationship between the modalities).

This whole-brain model successfully predicted EEG power in out-of-sample subjects for both alpha (‘brainwide’ vs ‘control’, Fig. 1d, *t* (23)=4.7, *p*<0.001) and delta (‘brainwide’ vs ‘control’, Fig. 1f, *t*(31)=14.8, *p*<0.001). Delta predictions surpassed the expected performance ceiling, displaying a mean correlation between predictions on held-out subjects and ground truth of *r*=0.41, and alpha was predicted at *r*=0.30. The models thus robustly predicted fluctuations of neural rhythms in held-out subjects, demonstrating the ability to learn dynamics shared across individuals. Representative predictions demonstrate that the model successfully captured short- and long-timescale fluctuations in alpha and delta power (Fig. 1e, 1g).

### Cortical, subcortical, and non-neural fMRI signals differentially predict alpha and delta rhythms

Given that we were able to predict neural rhythms on out-of-sample subjects, we next explored how we could use this approach to identify which regions carry information about alpha and delta fluctuations. Importantly, both neuronal and non-neuronal signals (such as large blood vessels and cerebrospinal fluid) carry information about general fluctuations in arousal state that are coupled to neural rhythms ^13^. We therefore first examined whether this predictive information was specific to gray matter regions in cortex or subcortex, or whether it was also carried in non-gray matter regions such as the white matter and the ventricles. We trained new models under three conditions: cortex only, subcortex only, and non-gray matter regions only, with testing again performed on held-out subjects (Fig. 2a). These conditions were then compared with the ‘brainwide’ and ‘control’ (fMRI data shuffled within each run) conditions described above.

**Fig. 2:**
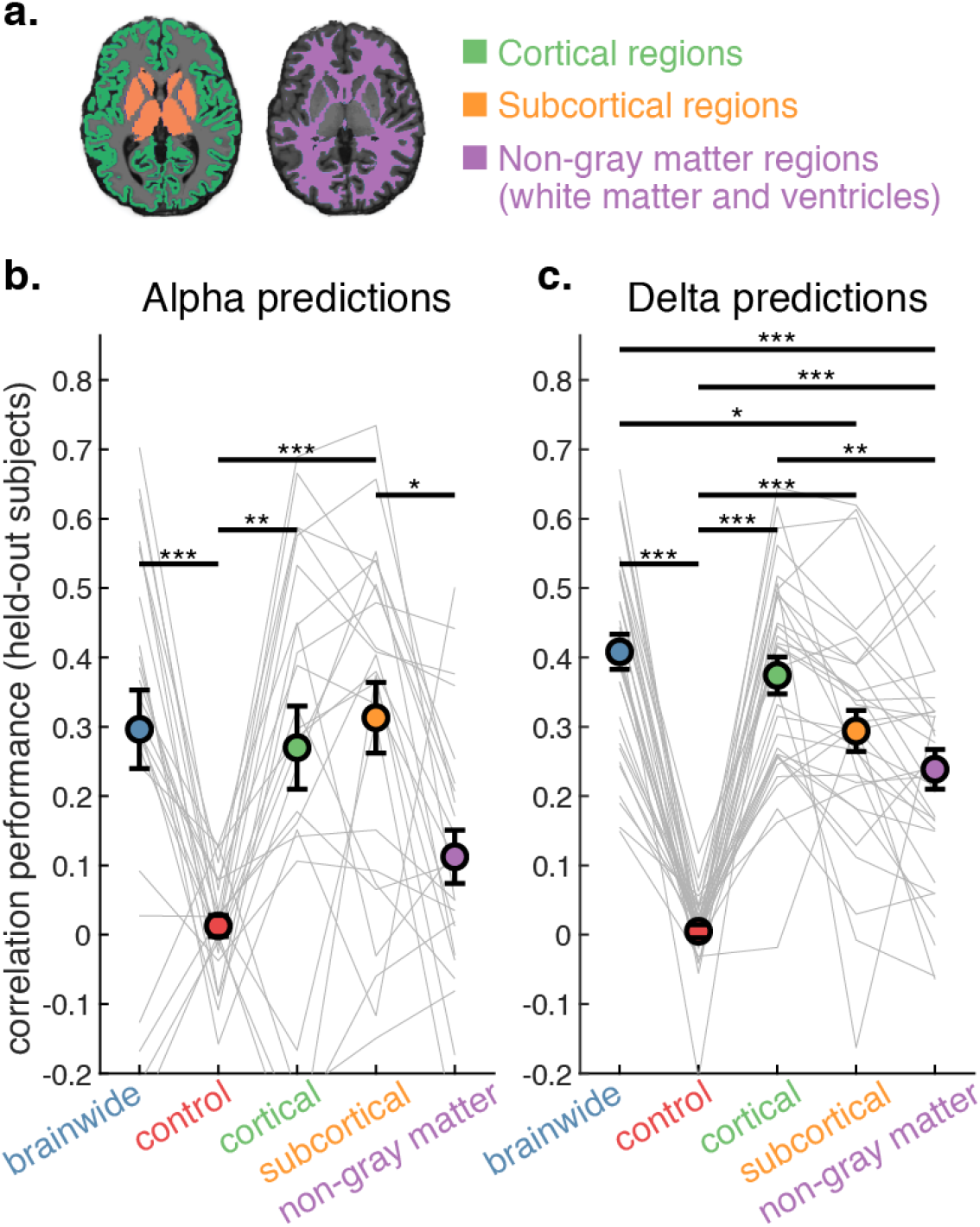
Alpha and delta rhythms show distinct relationships to cortical, subcortical, and non-neural fMRI signals. **a.** Models were trained and tested under three additional conditions: cortical regions only, subcortical regions only, and non-gray matter regions only (ventricles and global white matter). Performance was compared to models trained using all regions, and under a control condition using all regions but with shuffled fMRI data, breaking the true relationship between EEG and fMRI. **b.** Alpha power can be predicted by cortical and subcortical regions, but not by non-gray matter regions. Circles show mean correlation between alpha power ground truth and model predictions (on held-out subjects). * p<0.05; ** p<0.01; *** p<0.001; Tukey’s HSD. Error bars are SEM. Gray lines indicate individual held-out subject prediction performance. N=24. **b.** Delta power can not only be predicted by cortical and subcortical regions, but also by non-gray matter regions on their own. N=32.

Alpha power (Fig. 2b, Extended Data 1) was successfully predicted above chance by cortical (*r*=0.27, *p=*0.002) and subcortical (*r*=0.31, *p*<0.001) fMRI signals, with prediction performance achieving a similar level as the brainwide condition (*r*=0.30). The finding that subcortical regions alone could predict alpha rhythms on a held-out subject with such high accuracy was surprising, as these regions have a lower signal-to-noise ratio. Finally, alpha power could not be significantly predicted by the non-gray matter regions on their own, indicating that alpha-predictive fMRI activity relied on the specifically neural information present in the BOLD signal.

Delta power (Fig. 2c, Extended Data 2), on the other hand, could not only be predicted by neural sources, but was also significantly predicted (with worse performance) by the non-gray matter regions on their own (*r*=0.24, *p*<0.001), likely owing to the known coupling between delta power, cerebrospinal fluid flow (in the ventricles), and global blood flow ^13^. In other words, the delta-predictive fMRI signals possessed a significant component reflected outside of gray matter activity; however, the brainwide delta-predictive fMRI information also relied on specifically neural information, as the non-gray matter condition generated worse predictions than the brainwide condition (*r*=0.41, *p*<0.001) and the cortical condition (*r*=0.37, *p=*0.001). The cortical condition achieved similar predictive performance as the brainwide condition, but the subcortical condition performed significantly worse (*r*=0.29, *p=*0.01), indicating that delta predictions did not benefit as strongly from subcortical information as alpha did.

Given that our study employed two distinct datasets, collected at different sites, we also conducted statistical analyses on each dataset independently (Extended Data 3). The distinct predictive patterns for alpha and delta were consistent across the datasets.

### Individual fMRI regions can predict neural rhythms, with distinct spatial patterns for alpha and delta

To next identify finer-scale information beyond the coarse anatomical divisions of cortex and subcortex, we tested the information contained within each region in isolation by training and testing the model on a single bilateral gray matter region at a time. This prediction approach allows for the modeling of flexible temporal functions shared across subjects linking each neural rhythm to each fMRI area. We initially explored an ablation analysis, removing bilateral brain regions from the model and assessing the effects on performance, but found that performance changes were non-significant. Thus, we opted for training the model on individual bilateral regions instead.

We generated spatial maps of individual brain regions’ predictive performances for alpha and delta, to visualize the large-scale patterns of information content across the brain (Fig. 3). For visualization, the delta results combined the performance on held-out subjects for both datasets (*n=*32; model training was carried out independently in each dataset due to different imaging parameters). Alpha spatial maps included only Dataset A (*n=*18) as only 6 subjects in Dataset B displayed alpha rhythms, too few to assess predictions when using only a single region. These maps demonstrated that single brain regions could be sufficient to achieve successful prediction of neural rhythms, but with poorer performance than with a larger network. A control analysis (Extended Data 4) confirmed the significance of the global predictive maps by comparing its patterns to shuffled data.

**Fig. 3:**
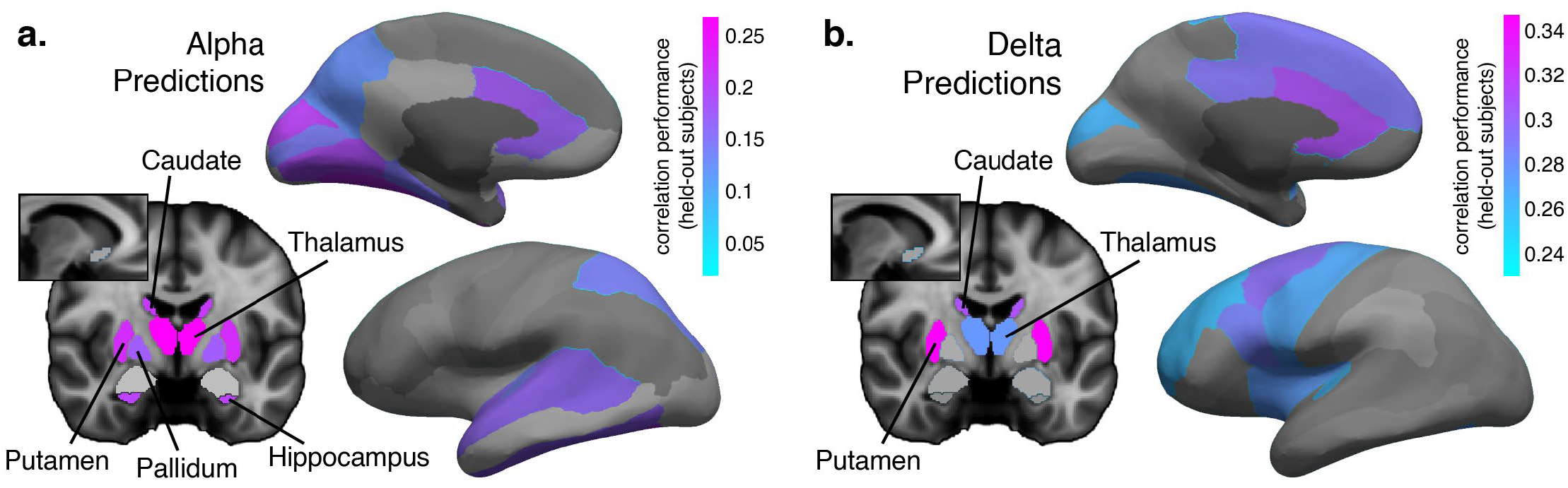
Prediction performance maps show which regions individually carry significant information about alpha and delta rhythms. **a**. Prediction performance (mean correlation between alpha predictions on held-out subjects and ground truth) when model was trained on single bilateral regions to predict alpha power. Regions shown in color yielded alpha predictions that were significantly better than control (non-significant regions are shown in grayscale). See Extended Data 5 for a list of correlation performance values. N=18. **b.** Prediction performance for delta. Model was trained on single bilateral regions combined with non-gray matter areas to predict delta power, in order to account for the delta-predictive power of the non-gray matter (see Fig. 2). Regions shown in color yielded predictions that were significantly better than non-gray matter areas alone. See Extended Data 6 for a list of correlation performance values. N=32.

This individual region analysis re-confirmed the predictive power of subcortical regions for alpha rhythms (Fig. 3a, Extended Data 5), revealing that alpha could be significantly predicted using only the thalamus (*r*=0.27, *p*<0.001), the dorsal striatum (separately as the putamen, *r*=0.23, *p=*0.002, and caudate, *r*=0.21, *p=*0.008), the hippocampus (*r=*0.20, *p=*0.002), or the pallidum (*r*=0.18, *p=*0.01). These results highlighted that neural rhythm prediction can be successful even in individual deep brain regions with lower signal-to-noise ratio. Consistent with the established coupling between alpha oscillations and fMRI activity within the visual system ^19,27,46–49^, cortical areas capable of individually predicting alpha power primarily comprised regions that process visual information (see Extended Data 5 for all correlation values and p-values), including early visual cortex (encompassing the cuneus, lingual, and pericalcarine), higher-order visual cognition regions (fusiform, inferior temporal, parahippocampal, superior parietal, precuneus), and multisensory regions (superior temporal and its bank, and transverse temporal ^50^). Alpha rhythms were also successfully predicted by the anterior cingulate cortex (separately as caudal and rostral), a region linked to arousal and reported to be correlated with thalamic activity ^51^. Prior EEG-fMRI studies identified correlations between alpha rhythms and the thalamus and occipital cortex ^19,27,46–49^; our results demonstrated that alpha fluctuations also reflect changes in neural activity in additional arousal-coupled cortical and subcortical regions (while the dorsal striatum and pallidum are not as commonly associated with arousal as the thalamus, they have demonstrated roles in controlling arousal; see ^52^ for a review), and to regions downstream of occipital cortex that receive visual information (such as the hippocampus), highlighting this predictive approach’s ability to learn more complex functions relating EEG and fMRI signals.

For the delta analysis (Fig. 3b), in order to identify delta-predictive information that is unique to gray matter regions and not reflected in systemic physiology, we accounted for the fact that delta power could be predicted from the non-gray matter regions alone (Fig. 2), by using this non-gray matter delta model as a control condition (in other words, we sought to identify region-specific information that is not also reflected in global physiological changes coupled to delta activity ^53,54^). While our preprocessing pipeline removed cardiac and respiratory cycles directly, low-frequency (<0.1 Hz) fluctuations due to slower physiological modulation (e.g. heart rate variability) remained ^55^, and these signals can vary substantially across arousal states ^56^. For each gray matter region, we trained a model that included both the region in question along with the non-gray matter regions, and then compared its performance to the model that used only the non-gray matter regions. This approach allowed us to avoid potential issues related to nuisance regressions ^57–60^. Delta fluctuations were predicted most strongly by the putamen (*r=*0.35, compared to the non-neural model, *r=*0.24; *p*<0.001), again demonstrating the predictive power of subcortical regions despite their lower signal-to-noise ratio. The caudate (*r=*0.31, *p=*0.001) and thalamus (*r=*0.28, *p=*0.04) also contained delta-predictive information significantly above the non-neural control (likely pointing to the shared role of arousal in predicting alpha and delta rhythms ^61^). The putamen and thalamus were previously associated with delta power in EEG-PET studies ^62,63^. The cortical delta-predictive information was distributed distinctly from alpha, favoring the anterior part of the brain (a prevalent initiating site for delta oscillations ^64^), including the anterior cingulate regions, which have previously been implicated in delta-band wave propagation ^64^.

### Two focused networks predict alpha, whereas delta information is diffuse

Our individual-region approach demonstrated that predictive information is present even on a single-region level, with distinct predictive maps for alpha and delta rhythms. These maps spanned a substantial portion of the cortex and subcortex, leading to the question of what manner of information about the neural rhythms is represented in each of those regions, and how this information may be separated into networks.

We therefore developed an analysis to identify predictive information shared across brain regions (Fig. 4a). We first assessed the predictive performance of each pairwise combination of fMRI regions in relation to the performance of its individual components, in order to evaluate whether regions contained distinct and complementary information (i.e., greater performance for the pair) or similar and redundant information (i.e., same or worse performance for the pair). In other words, we trained and tested the model on each pair of regions, and investigated the performance benefits (or losses) of each pair of regions (pair performance is shown in Extended Data 7), as compared to the individual regions performance (Fig. 3). As before, non-gray matter regions were also used when generating delta predictions, to identify gray matter-specific fMRI information beyond that present in the white matter or ventricles, as done in the prior analysis. Next, we performed k-means clustering on these results, using the gap statistic to evaluate the optimal clustering solution (including the null model of no suitable clustering) ^65^. By clustering the pairwise performance benefits, we could identify the presence of different networks of information: groups of regions that share redundant information among themselves (resulting in performance losses or no benefits when paired together), but which generate performance benefits when paired with regions from other predictive networks.

**Fig. 4:**
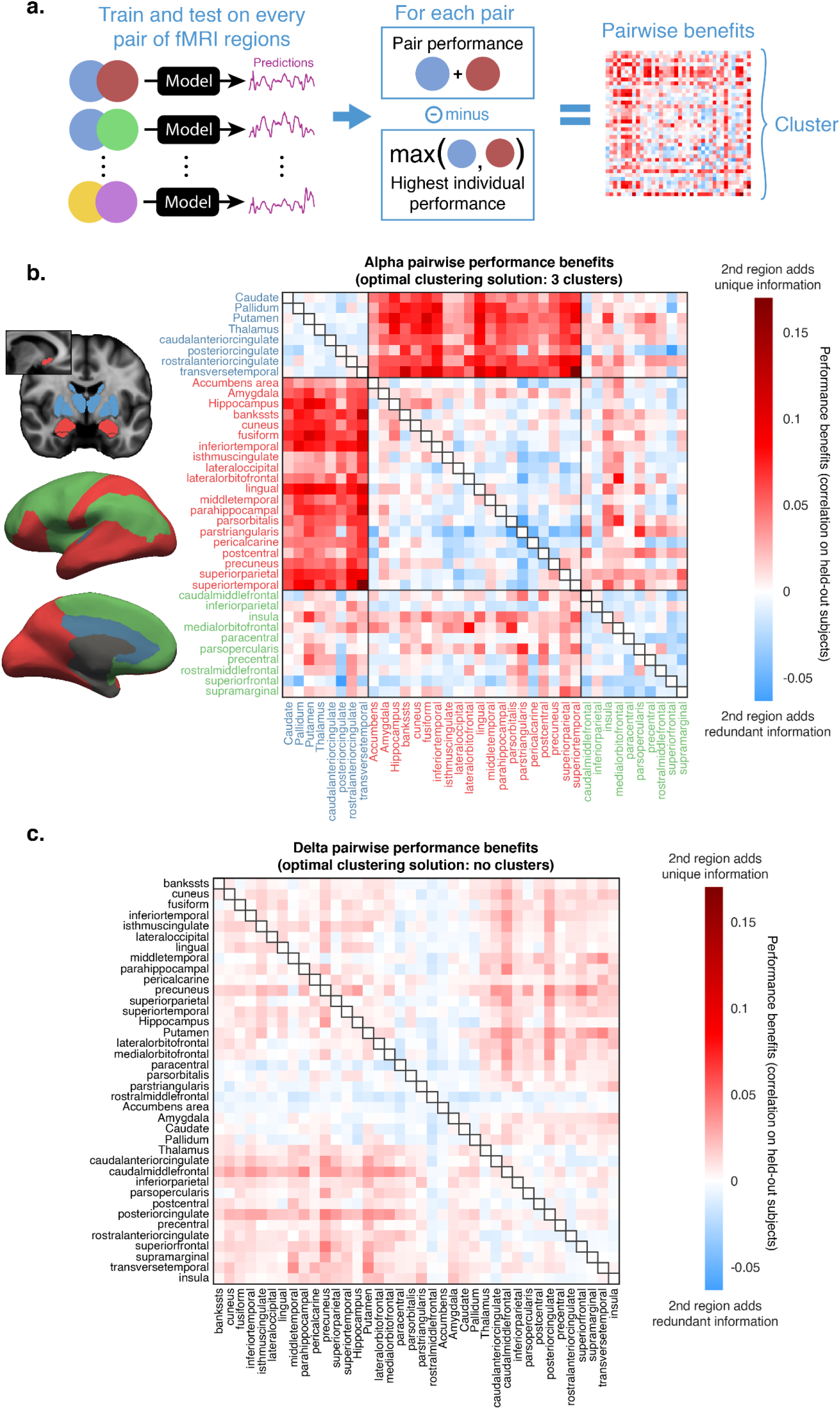
Clustering analysis of pairwise performance reveals two distinct networks of alpha-predictive fMRI information, while delta-predictive information is not separable in clusters. **a.** Models were trained on every possible pair of gray matter regions, and correlation performance was calculated on held-out subjects. Pairwise performance benefits were calculated as the performance gain (or loss) of a pair in relation to the maximum performance of its individual components; see Fig. 3 for individual performance values. Performance benefits were then clustered and evaluated with the gap statistic (Extended Data 8) to determine the optimal clustering solution. **b.** Identified clusters and all pairwise performance benefits for alpha-predictive information. When regions from the blue and red cluster were paired together, they consistently obtained performance benefits not found in any other cluster match. N=18. **c.** The clustering analysis revealed that delta did not possess a suitable clustering solution, as evidenced by no systematic pairwise performance benefits following a forced 3-cluster fitting. N=32.

We found that alpha pairwise performance benefits were optimally clustered in three groups (Extended Data 8). Two of these groups (Fig. 4b, blue and red clusters), when paired together, displayed substantial performance benefits (*r=*0.058, *95% CI* [0.054, 0.062]). Meanwhile, the third group (Fig. 4b, green cluster) included only regions that could not significantly predict alpha power (see Fig. 3a), and generated virtually no performance benefits when paired with either of the other groups (*r=*0.007, *95% CI* [0.002, 0.012]; *r=*0.008, *95% CI* [0.005, 0.011]). None of the clusters generated benefits when paired with themselves (*r*=−0.007, *95% CI* [−0.01, −0.003]; *r*=−0.004, *95% CI* [−0.007, −0.002]; *r*=−0.011, *95% CI* [−0.015, −0.007]). These results indicated that the blue and red clusters represented two distinct networks with different alpha-predictive information, while the green cluster represented regions which do not contain appreciable alpha-predictive information. A complementary analysis (Extended Data 9) found that correlation between the BOLD signals of different fMRI regions (as in typical functional connectivity analyses) did not capture the same marked clustering behavior, confirming that this analysis identified network structures beyond what can be captured from traditional fMRI approaches.

The smaller cluster of alpha-predictive information (Fig. 4b, blue) included the thalamus, the dorsal striatum (putamen and caudate), the pallidum, the anterior (separately as caudal and rostral) and posterior cingulate cortices, and the transverse temporal cortex. These results suggest that one component of alpha predictions reflected activity variance in this network of regions linked to arousal state modulation ^51,52^. The larger cluster of alpha-predictive information (Fig. 4b, red), included visual cortex (both the ventral and dorsal visual streams), prefrontal and temporal regions associated with higher-order visual cognition, and three deep-brain regions that are responsive to contents of visual information (the accumbens, amygdala and hippocampus), suggesting a distinct category of alpha-predictive information present in a network of regions involved in visual processing.

In contrast, the gap statistic demonstrated that the delta results did not present a suitable clustering solution (Extended Data 8). To illustrate the distinct network structures that represent alpha and delta, we forced a three-cluster fitting on the pairwise delta predictions (Fig. 4c), showing that delta did not display the clustering behavior present in the alpha analysis (Fig. 4b). Delta-predictive information, therefore, was not expressed as separable networks.

To obtain further insights on the network structure behind these rhythms, and particularly to better understand why delta did not possess a suitable clustering solution, we conducted an additional analysis to assess the size of fMRI networks required to predict each EEG rhythm. In other words, how many distinct fMRI regions are needed for optimal predictions? If many regions are required, this implies that predictive information is widely distributed across the brain, with low redundancy between regions. On the other hand, if only a small subset of regions is required for optimal predictions, this suggests that predictive information is very redundant across regions. To identify this number of optimal regions, we designed an iterative procedure to remove redundant information: in this procedure, low-weight regions are iteratively removed from the model (starting with all 84 regions) while performance changes are tracked.

We found that removing low-weight regions enabled us to achieve improved predictions of both alpha and delta power in relation to using brainwide data (Extended Data 10). These performance increases peaked at a larger number of regions for delta than for alpha (37 regions, vs. 18 regions), and subsequent reductions in region numbers caused performance to drop, eventually reverting the benefits of feature selection. This turning point happened only at very low numbers for alpha predictions (8 regions), indicating that alpha-predictive information has very low dimensionality. This finding was consistent with alpha possessing a small number of separable networks of shared information. Meanwhile, for delta predictions, the model degraded at a much larger number of regions (17 regions) with a more pronounced performance loss afterwards, indicating that unique delta information is present in a large number of regions with lower redundancy between regions. Finally, performance improvements at the optimal iteration were more pronounced for alpha (*r*=0.46 when using 18 regions, compared to brainwide *r*=0.30, *t*(17)=3.32, *p*=0.008) than delta (*r*=0.45 when using 37 regions, compared to brainwide *r*=0.41, *t*(31)=4.30, *p*<0.001), further pointing to alpha benefitting more strongly from feature selection due to the higher redundancy of its representation. This analysis also exemplifies why an ablation approach was not suitable for determining the unique contributions of each brain region, as both neural rhythms are represented by a substantial amount of redundant information across brain regions (with substantially more redundance for alpha).

Taken together, these analyses revealed two highly separable networks of alpha-predictive information, representing the dissociable components of arousal and visual dynamics, with a large amount of information redundancy within each network. Meanwhile, delta information was highly diffuse, requiring a large array of brain regions for optimal predictions.

## DISCUSSION

Neural rhythms signal changes in vigilance and play major roles in human cognition, yet the underlying network patterns they represent are difficult to investigate. We used machine learning to predict continuous fluctuations in EEG power in the alpha and delta bands from brainwide fMRI data, and achieved robust performance in held-out subjects, with a mean correlation value of *r*=0.41 and *r*=0.30 (respectively for delta and alpha; Fig. 1) between predictions and ground truth using brainwide data, and *r*=0.45 and *r*=0.46 following feature selection (Extended Data 10). This prediction-based approach demonstrated reproducibility across datasets (Extended Data 3), and was able to uncover information about continuous neural rhythm variations in individual fMRI regions (Fig. 3). We discovered that distinct network patterns were coupled to each rhythm, with strikingly differentiated spatial scales: two focused, separable networks predicted alpha rhythms from moment to moment, whereas a highly diffuse array of primarily cortical regions provided information about delta oscillations.

Alpha rhythms have been widely studied in the EEG-fMRI literature, and individual fMRI regions have been correlated to alpha power variations. Our findings replicate prior relationships between the thalamus/occipital cortex and alpha power ^18,19,24,27,46–49^, and identified additional relationships between alpha power and fMRI activity in several cortical and subcortical regions. Our results also demonstrated that separable networks are coupled to alpha rhythms: the dynamics within the visual system itself, and the modulation from the arousal system. Alpha rhythms have previously been correlated to activity in brain regions associated with vision and attention/arousal ^20,21,25,66,67^; our approach demonstrate that these networks contain highly separable information on the rise and fall of alpha power, and delineate which regions comprise each alpha-predictive network. The thalamus, which has often been viewed as the primary orchestrator of alpha rhythms ^19,24,47,48^, was revealed to be part of a circuit involving other key arousal regions: the dorsal striatum, the pallidum (which receives projections from the dorsal striatum, and in turn projects to the thalamus ^68^), and the cingulate cortex (which closely communicates with the thalamus ^69,70^). Meanwhile, distinct alpha-predictive information was present in the visual system (and in areas that respond to visual information), which may reflect that cortical neurons can intrinsically fire in the alpha range independent of thalamic input ^71^, and that alpha rhythms have been proposed to be involved in visual information flow between lower-level perceptual areas and higher-order cognitive regions ^15,72,73^.

We found a distinct network pattern linked to delta power, which indexes several sleep oscillatory dynamics, including slow oscillations and slow waves. While prior EEG-fMRI studies have identified fMRI activity locked to individual slow wave events ^74–76^, delta power variations represent distinct information, capturing continuous brain states as opposed to discrete neural events. Our work shows that delta predictions rely on combined information across a large number of fMRI regions, consistent with prior findings that correlations between delta power and fMRI are widespread across the cortex ^28,29^. Our approach revealed additional details about this widespread delta-coupled network: delta predictions did not form separable clusters, indicating that subtle differences in fMRI activity, primarily across the cortex, came together to predict variations in delta rhythms; this is also supported by the coarse-grained analysis (Fig. 2c) identifying superior delta predictions when using the full conjunction of cortical regions. Thus, delta predictions represented an accumulation of diffuse activity primarily distributed throughout the cortex. What dynamics within this broad network, with each region possessing a small amount of unique information, could be predicting delta power? Prior work has shown that arousal fluctuations are associated with cortical infra-slow propagating fMRI activity ^77^, and that infra-slow fMRI is coupled to delta power ^78^; it is therefore possible that this propagating infra-slow fMRI activity was core to the delta predictions. The varying phase across brain regions may explain why delta-predictive information required a large number of regions and was not separable in different networks. In addition, delta power itself exhibits complex propagating spatiotemporal dynamics ^79^, and can be both local and global in nature ^5,80^, suggesting that this diffuse fMRI predictive network may also represent a collection of diverse local slow wave dynamics. Future work could explore these questions by identifying traveling wave events in simultaneous EEG-fMRI and investigating whether distinct spatiotemporal features of infra-slow fMRI oscillations predict local or global delta power variations.

Delta predictions also substantially benefitted from information outside of the gray matter. While systemic physiology is often seen as noise in the context of fMRI, recent work has shown that these signals carry important information about brain state and thus neural activity ^53,54,56,60^. Our results indicate that delta rhythms are more strongly coupled to these systemic changes, reflected in the non-gray matter regions, than alpha rhythms. Our approach also provides a way to quantify information that is not reflected systemically, which may be advantageous for describing subtle changes in neural activity that are local, rather than global. However, in studies where the goal is to achieve the most accurate EEG predictions possible, preserving this systemic information could in turn benefit prediction accuracy.

We highlight that alpha power can be high in states other than wakefulness (such as during REM sleep, though only one subject displayed it in our data), and delta power can be high in states other than sleep; additionally, both of these bands display substantial variations during both sleep and wakefulness (see Fig. 1a), and our model was trained to capture these variations regardless of state. Thus, the networks we identified should not be interpreted as exclusively representing wake (and high alpha) or NREM sleep (and high delta); rather, they represent the continuous variations in these bands regardless of state. We particularly emphasize that NREM sleep is characterized by several additional frequency bands such as sigma (sleep spindles), which would be expected to display their own underlying network structures. Future work could attempt to investigate the underlying networks for other arousal-related frequency bands.

fMRI data is strongly characterized by network-level activity, with variations in multiple regions generating distinct spatiotemporal patterns ^81,82^. Our approach not only allows us to identify which regions are individually coupled to neural rhythms, but also to characterize whether the information across regions is unique or redundant. This provides a way to identify fMRI networks that are coupled to non-fMRI signals. As an example, we found that both the transverse temporal and superior temporal cortices, regions that are most commonly associated with auditory perception, were individually predictive of alpha rhythm variations to a similar degree. However, we found that they were in separate networks, with the superior temporal cortex—known to also play a role in visual perception ^50^—being in fact part of a network of visual alpha dynamics. Meanwhile, the transverse temporal cortex shared the same category of alpha-predictive information as an array of subcortical arousal-controlling regions, which may be explained by its activity reflecting thalamic-mediated state modulation, with prior work finding that the transverse temporal cortex is affected by sleep spindle density ^83^ and poor sleep quality ^84^.

Predictive machine learning has supported neuroscientific insights in event-related classification analyses across a range of neuroimaging modalities ^85–88^, and prior work has shown that some localized fMRI activity can be predicted from the EEG signal ^89,90^. Our method extended these predictive frameworks to achieve continuous, dynamic predictions of neural rhythms (which are much higher frequency than the fMRI temporal resolution) from brainwide fMRI data, in the absence of external stimuli and task conditions. This approach has several benefits, such as accounting for systemic signals in non-gray matter regions (by including the white matter and ventricles as predictors); ensuring that effects are generalizable across subjects through cross-validation; and allowing for the identification of multi-region relationships that represent shared or unique information. fMRI studies often use hemodynamic functional connectivity (defined by correlation patterns between different fMRI regions or voxels) to identify network dynamics; while this approach can identify broadly coherent neural activity, some aspects of electrophysiological networks are not captured in these correlation patterns, and electrocorticography work has shown that distinct neural frequency bands form spatially specific networks ^91^. Our results identified distinct fMRI networks for alpha and delta, providing a non-invasive way for identifying rhythm-specific networks. Finally, this predictive approach can be generalized to other continuous crossmodal analyses in neuroimaging data, to identify how spontaneous brain dynamics are coupled in a broad range of questions. Future work could identify distributed spatial dynamics underlying additional brain rhythms, investigate how these networks of information may vary across cognitive tasks, or probe how these networks may shift in aging and disease.

For this proof-of-concept, we opted for a relatively coarse anatomical parcellation as a means of dimensionality reduction, with 31 cortical and 7 subcortical parcels in each hemisphere, for a total of 76 gray matter parcels (activity within the voxels of each parcel were averaged). This averaging process provides dimensionality reduction and improved signal-to-noise ratio, and allows for bridging across different subject’s anatomies. Voxel-level activity was generally correlated with the mean signal in the parcels (Extended Data 11), and voxel-level variability did not appear to be meaningfully related to regional performance (see Fig. 3 and Extended Data 11), but several voxels exhibited distinct activity from the average, and this activity could contain additional predictive information about the neural rhythms. Future work using a more complex model could determine whether finer parcellation approaches, alternative methods of dimensionality reduction (such as principal component analysis), or even voxel-level predictions are capable of identifying dynamics beyond the benchmarks we established, which could allow for the identification of subtler information about neural rhythms. Voxel-level predictions in particular would be highly valuable if they could be achieved as they would enable network investigation with higher spatial precision; however, a substantial barrier is the high noise levels and high inter-subject variability when analyzing single voxels, and novel methodological approaches may be needed to achieve single-voxel-based predictions that can generalize to subjects outside of the training set.

In conclusion, we developed a novel approach to identify brainwide activity underlying fluctuations in neural rhythms, and demonstrated its results by investigating alpha and delta rhythms across sleep and wakefulness. Each rhythm was captured by distinct aspects of fMRI signals, with alpha predictions driven by two separable networks—one representing the visual system, and the other representing arousal-controlling regions—and delta being predicted by a diffuse array of primarily cortical regions. This work demonstrates that brainwide hemodynamics can predict EEG rhythms, and develops a framework for EEG prediction that could be used to study brain states in a range of cognitive tasks and populations.

## METHODS

### EEG-fMRI acquisition

Both datasets analyzed in this work were collected as part of other studies. Collection for Dataset A was ongoing while our analyses were conducted; the reported number of subjects for this dataset is the number that had been collected at the time our analyses began. Subsequently collected subjects were not counted or considered for the purposes of this study. Dataset B is from previously published studies ^13,51^.

All subjects reported no neurological, psychiatric, or sleep disorders, and were screened for the following: no MR contraindications, have habitual sleep duration of no less than six hours, daily caffeine consumption of less than 300 mg, not smoke cigarettes, weigh less than 250 pounds, not be pregnant, be comfortable sleeping on their back, and not be taking medications that affect sleep.

Subjects were instructed to lie still inside the scanner with their eyes closed and to allow themselves to fall asleep. In Dataset A, eye closure was asserted with eye tracking. A subset of subjects (17 in Dataset A, 5 in Dataset B) was additionally instructed to press a button on every breath in and out, and to return to performing the task if they awoke during the scan; a lack of button presses indicated subject was asleep ^51^. Button press data was not analyzed in this study.

#### Dataset A

We obtained written informed consent from 22 healthy young adults; 21 (9 male, 12 female; mean age 23.4) were analyzed for the purposes of this study. One subject was excluded due to unusable EEG data caused by excessive subject motion. The protocol was approved by the Boston University Charles River Campus Institutional Review Board (where 17 subjects were collected) and by the Massachusetts Institute of Technology Institutional Review Board (where 5 subjects were collected). Subjects were scanned on a 3T Siemens Prisma scanner with a 64-channel head-and-neck coil. Anatomical references were acquired using a 1 mm isotropic T1-weighted multi-echo MPRAGE ^92^. Functional runs were acquired using TR = 0.378 s, TE = 31 ms, 2.5 mm isotropic voxels, 40 slices, Multiband factor=8, blipped CAIPI shift = FOV/4, flip angle=37°, no in-plane acceleration. Up to three successive runs of 25 minutes each were obtained per subject contingent on subject comfort.

EEG was acquired using MR-compatible 32-channel EEG caps fitted with 4 carbon wire loops (BrainProducts GmbH, Germany) at a sampling rate of 5000 Hz. EEG acquisition was synchronized to the scanner 10 MHz clock to reduce aliasing of high-frequency gradient artifacts. Additional sensors were used to record systemic physiology: respiration was measured simultaneously using an MRI-safe pneumatic respiration transducer belt around the abdomen and pulse was measured with a photoplethysmogram (PPG) transducer (BIOPAC Systems, Inc., Goleta, CA, USA). These systemic physiology signals were not analyzed in this study.

#### Dataset B

We used a previously published dataset which had collected data from 13 healthy young adults; 11 were analyzed for the purposes of this study (2 male, 9 female, mean age 26.7). One subject was excluded due to a software error that caused fMRI data to be collected at a different temporal resolution, and another was excluded due to poor-quality EEG data caused by excessive subject motion. The protocol was approved by the Massachusetts General Hospital Institutional Review Board. Subjects were scanned on a 3T Siemens Prisma scanner with a 64-channel head-and-neck coil. Anatomical references were acquired using a 1 mm isotropic T1-weighted multi-echo MPRAGE ^92^. Functional runs acquired 40 interleaved BOLD-weighted EPI slices with 2.5 mm^3^ isotropic voxels. fMRI protocols consisted of a single-shot gradient echo SMS-EPI ^93^ with MultiBand facto*r*=8, matrix=92×92, blipped CAIPI shift=FOV/4, T*R*=367 ms, TE=30-32 ms (changed midway through the study due to a software upgrade) nominal echo-spacing=0.53 ms, flip angle=32-37°, and no in-plane acceleration. VERSE factor was set between 1 and 1.5 depending on individual subject SAR constraints. Individual runs could last up to 2 hours. If runs ended earlier, subsequent runs would be started up to a maximum total scan duration of 2.5 hours as long as subjects were still comfortable and sleeping.

EEG was acquired using MR-compatible 256-channel geodesic nets and a NA410 amplifier (Electrical Geodesics, Inc., Eugene, OR USA) at a sampling rate of 1000 Hz. EEG acquisition was synchronized to the scanner 10 MHz clock to reduce aliasing of high-frequency gradient artifacts. The scanner cryopump was temporarily shut off during EEG acquisition to reduce vibrational artifact. To acquire reference signals to be used for EEG noise removal ^94^, subjects wore a reference layer cap composed of an isolating vinyl layer and conductive satin layer on the head, with grommets inserted to allow electrodes to pass through and make contact with the scalp, while other electrodes remained isolated from the scalp and recorded the noise, resulting in a total of 30–36 EEG electrodes per subject. Physiological signals were simultaneously acquired using a Physio16 device (Electrical Geodesics, Inc., Eugene, OR USA). ECG was measured through two disposable electrodes placed on the chest diagonally across the heart, with an MR-compatible lead (InVivo Corp, Philips). Respiration was measured through a piezoelectrical belt (UFI systems, Morro Bay, CA USA) around the chest.

### EEG pre-processing

#### Dataset A

Gradient artifacts were removed using average artifact subtraction ^95^, with moving average of the previous 20 TRs. Ballistocardiogram artifacts were removed from each EEG channel using signals from the 4 carbon wire loops using the sliding Hanning window regression method from the EEGlab CWL toolbox ^96^ with the following parameters: window = 25 s and a lag = 0.09 ms. Clean EEG signals were re-referenced to the average of EEG channels.

#### Dataset B

Gradient artifacts were removed using average artifact subtraction ^95^, with moving average of the previous 20 TRs. Electrodes were then re-referenced to the common average, computing this separately for electrodes contacting the head, and those placed on the reference layer. Channels on the cheeks and borders of the reference cap were excluded from the common average. Ballistocardiogram artifacts were removed using regression of reference signals from the isolated EEG electrodes ^97^. Since there was a larger number of noise electrodes than signal electrodes, the regression was performed after subsampling the noise electrodes, using only every fourth isolated electrode. Because the position of and physiological noise influences on the electrodes can vary over the long recording times used here, we implemented a dynamic time-varying regression of the reference signals. Beta coefficients for the best-fit regression within 30 s sliding time windows were fit using least-squares; these beta values were then linearly interpolated over the nonoverlapping windows. The resulting interpolated beta value at every time point was then used for a local subtraction of the reference signals from the modeled EEG recording. This regression was performed individually for each EEG channel.

### fMRI pre-processing

The cortical surface was reconstructed from the MEMPRAGE volume using Freesurfer version 6 ^98^. All functional runs were slice-time corrected using FSL version 6 (slicetimer; https://fsl.fmrib.ox.ac.uk/fsl/fslwiki) ^99^ and motion corrected to the middle frame using AFNI (3dvolreg; https://afni.nimh.nih.gov/). Each motion-corrected run was then registered to the anatomical data using boundary-based registration (bbregister) ^100^.

For Dataset A, physiological noise was removed using HRAN, open-source software implementing a statistical model of harmonic regression with autoregressive noise ^101^.

For Dataset B, physiological noise removal was performed using dynamic regression based on the concept of RETROICOR ^102^ and adapted for fast fMRI as follows. The respiratory trace was bandpass filtered between 0.16–0.4 Hz using a finite impulse response filter and the instantaneous phase was computed as the angle of the Hilbert transform. The cardiac peaks were detected automatically using the FASST toolbox (montefiore.ulg.ac.be/~phillips/FASST) and the phase was modeled as varying linearly between each identified peak. Sine and cosine basis functions using the phase of the signal and its second harmonic were generated as regressors for physiological noise. This regression was performed over 1000 s windows sliding every 400 s to enable high-quality physiological noise removal as the heart rate and respiratory rate varied throughout the scan. No spatial smoothing was applied.

We highlight that our physiological noise correction removed noise associated with specific cardiac and respiratory cycles, but did not regress out the low-frequency component of cardiac and respiratory rate variations, as regressing these low-frequency signals can also remove information about neural activity that is coupled to systemic physiology ^53,54^. Since our fMRI acquisition used very short TRs, these individual cardiac and respiratory cycles are not aliased in the data, and they can distort the BOLD signal with motion artifacts.

### Data preparation

Data preparation, model training, and statistical analyses were conducted in MATLAB unless noted otherwise. EEG power was extracted from occipital electrodes via the multitaper spectrogram method ^103,104^. We used occipital electrodes as they display less MR noise and improved data quality due to being pressed against the back of subjects’ heads as they lie down in the scanner. Spectrograms were obtained using a 5 second window, sliding every 0.1 s. Spectrograms were visually inspected for the presence of clear eyes-closed alpha rhythms. Most healthy adults display a clear increase in alpha power when awake with eyes closed (see Fig. 1a), but a small percentage of individuals do not show this increase for reasons not yet understood ^42–45^. As this difference may reflect distinct underlying networks, we excluded these subjects from the alpha training and analyses (3 subject in Dataset A, 5 subjects in Dataset B). These subjects were included in the delta training and analyses.

The power time series was then extracted for the frequencies of interest: 1.1-4 Hz for delta (chosen to minimize contamination from low-frequency artifacts in EEG-fMRI) and 8.5-12 Hz for alpha (chosen to minimize contamination from theta and spindle frequency ranges while still covering subject variations in alpha range). Power in the 40-52 Hz range was also extracted for the purpose of artifact detection (see below).

For dataset A, power was extracted from three occipital electrodes (O1, O2, and Oz) and then averaged. For dataset B, which used a reference layer for artifact removal ^94^ and therefore did not have identical channel layouts across subjects, power was extracted from the single occipital electrode closest to Oz which had good data quality based on visual inspection of the spectrogram. This inspection procedure was done for a prior study ^13^ and we used the same previously selected electrodes. Power time series were then interpolated to match fMRI TR timings.

fMRI parcellation was performed in FreeSurfer based on the Desikan-Killiany atlas ^105^. The following region labels were obtained: 31 bilateral cortical regions (the entire cortex minus the temporal pole, frontal pole, and entorhinal regions, which were not covered by the acquisition volume in all subjects), 7 bilateral subcortical regions (thalamus, pallidum, amygdala, caudate, putamen, accumbens, hippocampus), and 8 non-grey matter regions (left- and right-total white matter as an index of global blood flow, left- and right-lateral and inferior lateral ventricles, third ventricle, and fourth ventricle); total of 84 regions. Anatomical labels were interpolated to functional space, and functional voxels with 70% or above of overlap with the label were averaged to generate the mean region time series. The first 20 TRs of each run were discarded, and the averaged fMRI timeseries of each region (separately for each subject) was z-scored.

To mark data points unsuitable for analyses, the following criteria were used: 1) TRs with motion above 0.3 mm; 2) z-scored EEG power in the 40-52Hz range with a value above 1 (adjusted to be lower for runs with particularly noisy data); 3) detected local outliers (distance larger than 6 local scaled median absolute deviation (MAD)) within a sliding window of 1000 TRs (for dataset B, a threshold of 3 MAD was used instead due to noisier data; thresholds were also stricter for subjects with excessively noisy data). The outlier detection method was applied to delta power and power in the 40-52Hz range (but not to alpha power, as its naturally fast fluctuations would be erroneously detected as outliers; however, three subjects in Dataset A with excessively noisy data underwent alpha band outlier removal based on a smaller window). Datapoints that fulfilled any of these criteria were then removed, and the remaining data was saved as separate segments to avoid creation of spurious continuities (segments below 60 TRs were discarded, as 60 TRs of continuous fMRI are required to predict each EEG point). EEG data for each subject was then rescaled to 0-1.

### Model training and testing

Following the above, fMRI data was split into overlapping 60-TR sections, with the goal of predicting the single EEG power point at the center of the fMRI window (equivalent to the position of the 30^th^ fMRI point). We predicted the EEG point at the center of the window to avoid assumptions about temporal relationship between the two modalities (such as regions which might have early or late responses), as our goal in this work was to uncover spatial patterns. Early model explorations revealed similar prediction performance for segment sizes in the 40 to 80 TR range (roughly 14s to 29s); 60 TRs was ultimately chosen for being long enough to include the width of the canonical HRF in the post-EEG half of the segment, while being short enough to avoid excessively inflating the number of weights that the model had to learn.

In order to predict every EEG point that had been interpolated to match the fMRI TR timings, there was a 59-TR overlap between each adjacent predictor window (windows were slid by 1 TR). Given that no fMRI points would be available for EEG points at the edges of the data segments following artifact removal, the first 29 and the last 30 EEG points of each segment were discarded.

The model was trained on a maximum of 5040 predictors (60 TR segments and 84 parcellated regions; region list under ‘Data preparation’). Given that different analyses used different numbers of parcellated regions, the actual number of predictors varied by analysis (i.e. for the results presented in Fig. 3a, the model was trained on each bilateral region at a time; thus, only 120 predictors).

At each training fold, all data for one subject was held-out as test data. The linear regression model, specifically MATLAB’s *fitrlinear* function with the ‘leastsquares’ learner ^106^, was then trained on the remaining subjects. The function fits a beta weight value for each input predictor feature (each TR of each input fMRI region), along with a bias value, and a lambda regularization term. The regularization term multiples the sum of squares of all model weights to penalize large weights and prevent the model from overfitting (L2 regularization). During training, the model adjusts its weights and bias using stochastic gradient descent. We explored using more complex models, including neural networks with recurrent layers and temporal convolutions, but found that performance was not significantly better. Thus, we moved ahead with the simpler model.

In each fold, the lambda regularization term was adjusted through Bayesian hyperparameter optimization by partitioning the training data (i.e. the held-out subject was not used) in 5 folds over 30 iterations, with repartitioning at every iteration. The best lambda value was then selected, and the model was trained on all training data at once. While the model frequently chose very similar lambda values in each held-out subject fold, this additional computational effort resulted in a strict training regimen where no data from the held-out subject was ever used to tune the model. In earlier explorations of the model, the Bayesian hyperparameter optimization was also allowed to select between L1 and L2 regularization, and between least-squares and SVM learners, but given that L2 consistently outperformed L1, and least-squares consistently outperformed SVM, we fixed L2 and least-squares as options to reduce computational processing time.

Once the optimal lambda value was determined and the weights and biases were fit on the training data, the model was fixed and used to generate predictions on the held-out subject. We highlight that normalization (see ‘Data preparation’ above) was done separately within each subject so that at no point did the held-out subject data affect training data. A Pearson’s linear correlation coefficient was then calculated between the predictions made for the held-out subject and the ground truth.

In order to achieve more precise estimates of predictive performance, we repeated each train-and-test fold 10 times with distinct random seeds, yielding 10 trained models and thus 10 sets of predictions (plus 10 Pearson’s correlation values) for each held-out subject. In all analyses, each correlation value reported for each held-out subject represents the mean correlation performance of 10 models.

All model training was done separately on each dataset due to the usage of distinct imaging parameters (particularly the distinct TRs). After model training and testing on held-out subjects for each dataset, the performance values for held-out subjects was combined for statistics and visualizations.

### Statistical analyses

#### Major Anatomical groups analysis

The model underwent the training and testing routine described above using different groups of fMRI regions as input: only the subcortical regions, only the cortical regions, only the non-gray matter regions, and all of them together. Finally, the model also went through the same training routine using all the regions as input, but with fMRI data shuffled in relation to the EEG (which was kept unshuffled in order to preserve its original temporal structure). Once the correlation performance was obtained for all subjects and conditions, an ANOVA was conducted, with subject ID as a random variable, and condition as a fixed variable. Pairwise comparisons between each condition were then performed using Tukey’s HSD (honestly significant difference, two-sided), conducted within each dataset (A/B/combined) and analyses type (alpha/delta). Tukey’s HSD corrects for multiple comparisons and operates upon the ANOVA results ^107^.

#### Bootstrapping for single-region analysis

To assess whether prediction performance differences between individual regions were not the result of the ordering of random noise, we shuffled the region labels within each subject, then obtained region means, ordered the means, and fit a line to the ordered means. This was repeated 1000 times; the resulting lines were then averaged, and a 95% confidence interval (based on number of subjects) was calculated. Verification that the line fit to the real (non-shuffled) data was outside these bounds was taken to mean that region differences (i.e. spatial pattern of predictive performance across fMRI regions) were significant and not due to ordering of random noise. Following this analysis, paired two-tailed t-tests were conducted to assess which regions were superior to shuffled control (or to non-gray matter regions in the case of the delta analyses).

#### Clustering of pairwise performance

The model was trained and tested on each pair of gray matter regions. For delta, the non-gray matter regions were also added to each gray matter pair. For each pair, the performance values from the single-region analysis (correlation between predictions and ground truth) were noted, and the highest performance was subtracted from the pair’s performance. This value represented the performance gain (or loss) of the pair in relation to its best individual component. These values were then used for clustering.

To assess the optimal clustering solution (including the null hypothesis of no suitable clustering), we used the gap statistic ^65^. The gap value is defined as the difference between the within-cluster dispersion obtained by Monte Carlo sampling from a reference distribution (generated from a uniform distribution over a box aligned with the principal components of the data) and the actual within-cluster dispersion obtained by clustering the real data. The ideal number of clusters is defined as the smallest number of clusters that satisfies *Gap(k)*≥*Gap(k+1)*−*SE(k+1)*, where *Gap(k)* and *Gap(k+1)* are the gap values for the clustering solutions with k and k+1 clusters respectively, and *SE(k+1)* is the standard error of the clustering solution with k+1 clusters.

#### Spatial scales analysis

After training and testing of the model using all 84 parcellated regions, the model’s beta weights were extracted, and weights (across all TRs) for each region were squared and summed. The regions that displayed weights whose sum of squares were below the bottom 10% (as distribution percentiles) were then removed from the model, which was then once again trained and tested. This was repeated up until 5 regions were left in the model. Performance in each iteration was then compared to performance using all brain regions with two-tailed t-tests corrected using the Benjamini-Hochberg procedure ^108^.

## ACKNOWLEDGEMENTS

This study was funded by National Institutes of Health grants R01-AG070135, U19-NS128613, and U19-NS123717; the Sloan Fellowship, the McKnight Scholar Award, the Pew Biomedical Scholar Award, the Simons Collaboration on Plasticity in the Aging Brain (811231), and the One Mind Rising Star Award. Resources were provided by NSF Major Research Instrumentation grant BCS-1625552. Collaboration was facilitated by the Janelia Visiting Scientist Program. The funders had no role in study design, data collection and analysis, decision to publish, or preparation of the manuscript.

## Author contributions

L.P.L.J. and L.D.L. conceptualized the study. S.M.B, S.D.W., and L.D.L. designed the experiments. S.M.B and S.D.W. collected and pre-processed the data, and wrote the corresponding sections of the methods. L.P.L.J. conducted data preparation, performed the analyses, created the figures, and wrote the manuscript. C.S. advised on modeling and analytical methods. C.S. and L.D.L. revised the manuscript and all authors provided edits.

## LIST OF LEGENDS FOR SUPPORTING INFORMATION FILE

Extended Data 1: Tukey’s HSD statistical results for alpha predictions using major anatomical regions.

Extended Data 2: Tukey’s HSD statistical results for delta predictions using major anatomical regions.

Extended Data 3: Predictive patterns are consistent across datasets.

Extended Data 4: Single-region prediction performance patterns cannot be explained by the ordering of noise. Extended Data 5: Paired t-test results for regional alpha-predictive performance.

Extended Data 6: Paired t-test results for regional delta-predictive performance.

Extended Data 7: Distinct prediction patterns when model is trained on every possible combination of bilateral gray matter regions.

Extended Data 8: Clustering analysis reveals the optimal number of clusters that explain pairwise performance benefits for each frequency band.

Extended Data 9: Decoded information results in highly discrete networks (see Fig. 4), and only weakly reflects correlation between the BOLD activity of different brain regions.

Extended Data 10: Alpha rhythms can be predicted from a small spatial network, whereas delta requires information from a broader set of regions.

Extended Data 11: Correlation between individual voxels within a region, and the regional mean timeseries, across all subjects and runs.

