## Supplementary figures for "Brainwide hemodynamics predict EEG neural rhythms across sleep and wakefulness in humans"

| Group A | Group B | Lower Limit | Difference Mean | Upper Limit | p-value | q-stat |
|---|---|---|---|---|---|---|
| all | control | 0.099 | 0.283 | 0.468 | <0.001 | 6.009 |
| all | cortical | -0.158 | 0.026 | 0.211 | 0.995 | 0.560 |
| all | subcortical | -0.202 | -0.017 | 0.168 | 0.999 | 0.354 |
| all | non-gray matter | -0.001 | 0.184 | 0.369 | 0.052 | 3.896 |
| control | cortical | -0.442 | -0.257 | -0.072 | 0.002 | 5.449 |
| control | subcortical | -0.485 | -0.300 | -0.115 | <0.001 | 6.363 |
| control | non-gray matter | -0.285 | -0.100 | 0.085 | 0.568 | 2.113 |
| cortical | subcortical | -0.228 | -0.043 | 0.142 | 0.967 | 0.914 |
| cortical | non-gray matter | -0.028 | 0.157 | 0.342 | 0.134 | 3.336 |
| subcortical | non-gray matter | 0.016 | 0.200 | 0.385 | 0.026 | 4.250 |

***Extended Data 1: Tukey's HSD statistical results for alpha predictions using major anatomical regions*** *(Fig. 2b). An ANOVA first identified a main effect of model input condition on correlation performance (F(115)=7.8, p<0.001), and a Tukey's HSD analysis was then carried out to identify pairwise differences of correlation performance between the conditions. Lower and upper limits refer to the confidence interval of the pairwise difference.*

| Group A | Group B | Lower Limit | Difference Mean | Upper Limit | p-value | q-stat |
|---|---|---|---|---|---|---|
| all | control | 0.306 | 0.403 | 0.500 | <0.001 | 16.055 |
| all | cortical | -0.063 | 0.034 | 0.131 | 0.875 | 1.352 |
| all | subcortical | 0.017 | 0.114 | 0.211 | 0.011 | 4.548 |
| all | non-gray matter | 0.072 | 0.169 | 0.266 | <0.001 | 6.744 |
| control | cortical | -0.466 | -0.369 | -0.272 | <0.001 | 14.702 |
| control | subcortical | -0.386 | -0.289 | -0.192 | <0.001 | 11.507 |
| control | non-gray matter | -0.331 | -0.234 | -0.137 | <0.001 | 9.311 |
| cortical | subcortical | -0.017 | 0.080 | 0.177 | 0.158 | 3.195 |
| cortical | non-gray matter | 0.039 | 0.135 | 0.232 | 0.001 | 5.391 |
| subcortical | non-gray matter | -0.042 | 0.055 | 0.152 | 0.528 | 2.196 |

***Extended Data 2: Tukey's HSD statistical results for delta predictions using major anatomical regions*** *(Fig. 2c). An ANOVA first identified a main effect of model input condition on correlation performance (F(155)=40.3, p<0.001), and a Tukey's HSD analysis was then carried out to identify pairwise differences of correlation performance between the conditions. Lower and upper limits refer to the confidence interval of the pairwise difference.*

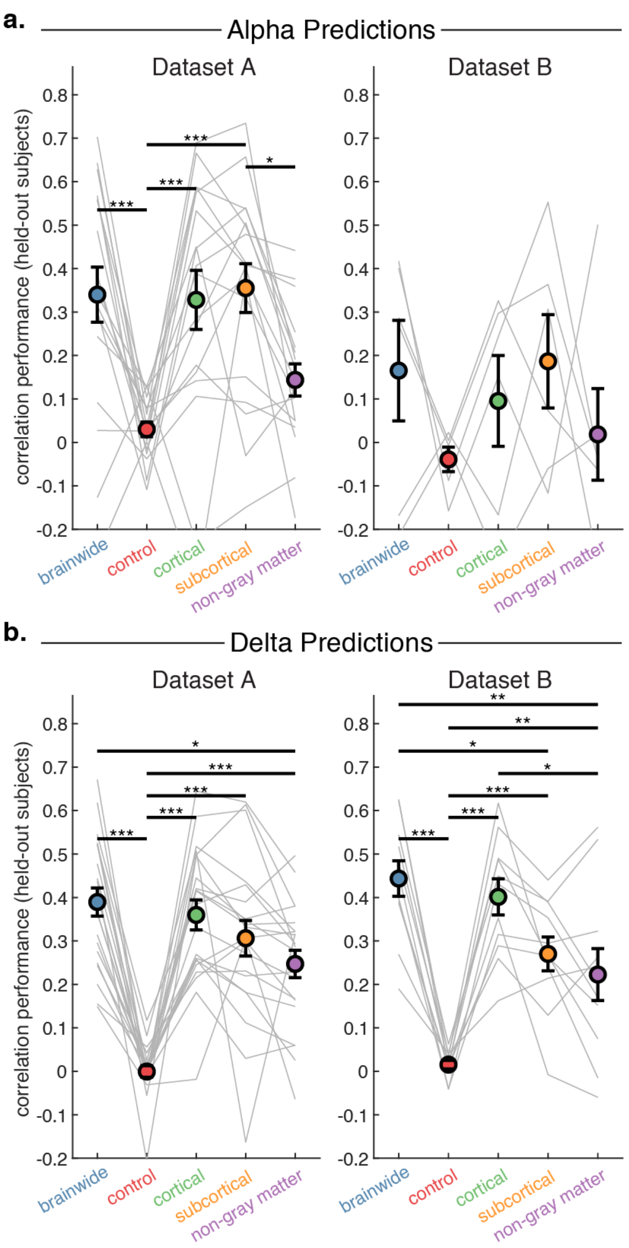


***Extended Data 3: Predictive patterns are consistent across datasets. a.*** *Alpha power prediction patterns favor subcortical regions, and alpha cannot be predicted from non-gray matter regions. Due to a low number of subjects (n=6) with alpha rhythms in Dataset B, results were not significant when examining this dataset on its own.* ***b.*** *Delta power prediction patterns favor cortical regions, and delta can be predicted from non-gray matter regions in both datasets. Circles show mean correlation between EEG power ground truth and model predictions (on held-out subjects). * p<0.05; ** p<0.01; *** p<0.001; Tukey's HSD. Error bars are SEM. Gray lines indicate individual held-out subject prediction performance.*

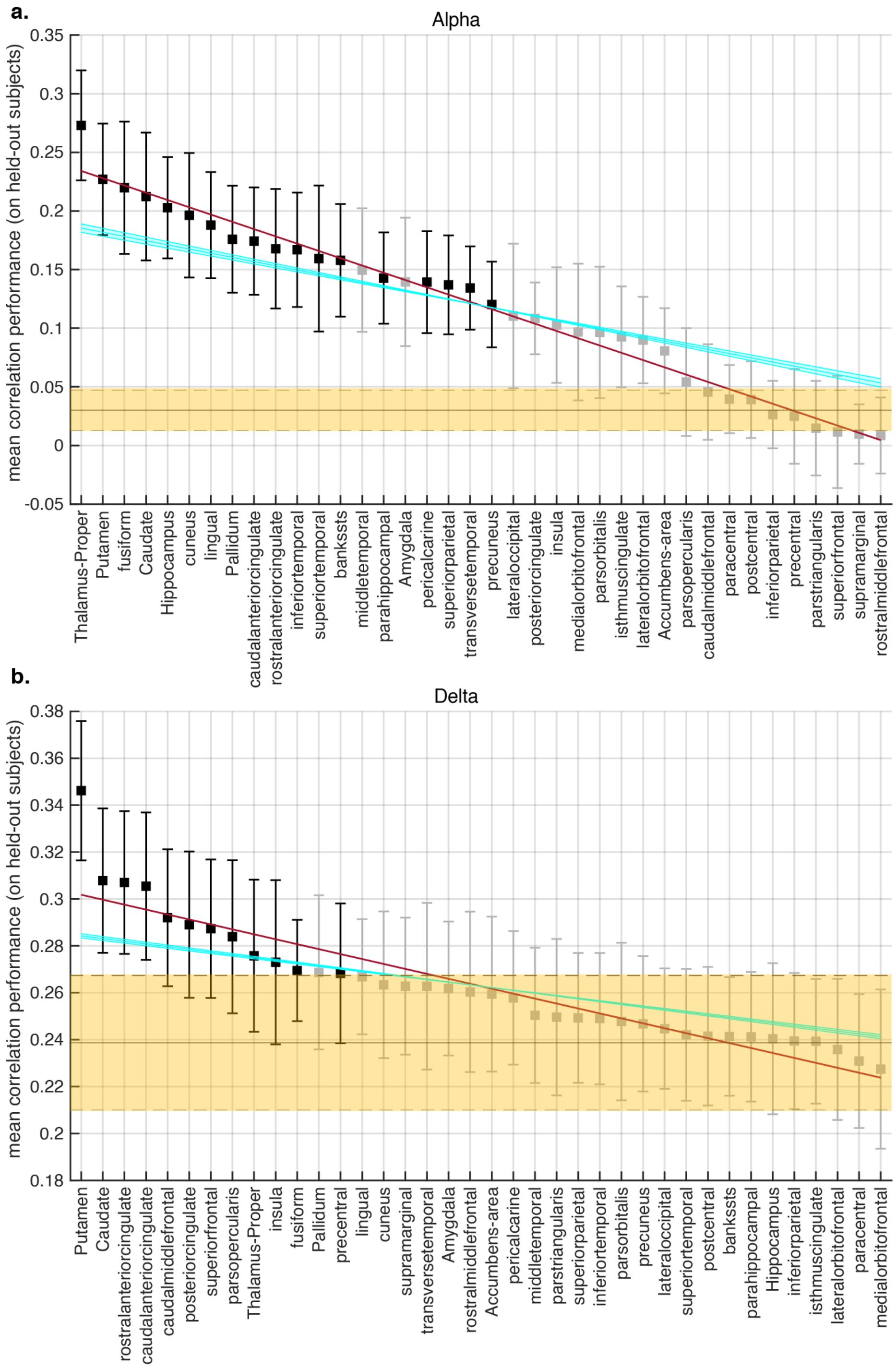


***Extended Data 4: Single-region prediction performance patterns cannot be explained by the ordering of noise.*** *A control analysis where region labels were shuffled (cyan lines with 95% confidence intervals, see methods for details) was contrasted with lines fitted to the actual results (red lines); the red lines display a steeper slope and are outside the confidence interval of the cyan lines, indicating that regional performance differences were not the result of the ordering of noise.* ***a.*** *Mean alpha-predictive performance of each fMRI region. The yellow area (mean with SEM) indicates the 'control' condition, which had the fMRI data shuffled in relationship to the EEG; regions plotted in black demonstrated performance significantly better than this condition, indicating they predicted alpha rhythms better than chance. Error bars are SEM.* ***b****. Mean delta-predictive performance of each fMRI region. The non-gray matter regions were included along with each bilateral gray matter region (see main text for justification). The yellow area (mean with SEM) indicates the 'non-gray matter' condition; regions plotted in black demonstrated performance significantly better than the non-gray matter regions, indicating that they contained delta-predictive information beyond what is present in non-neural sources.*

| Region | r | p-value | lower limit | upper limit | df | t-value |
|---|---|---|---|---|---|---|
| Thalamus | 0.273 | 0.0004 | 0.125 | 0.360 | 17 | 4.36 |
| Putamen | 0.227 | 0.0024 | 0.081 | 0.313 | 17 | 3.57 |
| fusiform | 0.220 | 0.0042 | 0.069 | 0.311 | 17 | 3.30 |
| Caudate | 0.212 | 0.0085 | 0.053 | 0.311 | 17 | 2.98 |
| Hippocampus | 0.203 | 0.0022 | 0.072 | 0.273 | 17 | 3.61 |
| cuneus | 0.196 | 0.0114 | 0.043 | 0.290 | 17 | 2.84 |
| lingual | 0.188 | 0.0040 | 0.058 | 0.258 | 17 | 3.32 |
| Pallidum | 0.176 | 0.0122 | 0.036 | 0.255 | 17 | 2.80 |
| caudalanteriorcingulate | 0.174 | 0.0146 | 0.032 | 0.256 | 17 | 2.72 |
| rostralanteriorcingulate | 0.168 | 0.0284 | 0.016 | 0.259 | 17 | 2.40 |
| inferiortemporal | 0.167 | 0.0172 | 0.027 | 0.246 | 17 | 2.64 |
| superiortemporal | 0.159 | 0.0490 | 0.001 | 0.258 | 17 | 2.12 |
| bankssts | 0.158 | 0.0192 | 0.024 | 0.232 | 17 | 2.59 |
| middletemporal | 0.150 | 0.0511 | -0.001 | 0.239 | 17 | 2.10 |
| parahippocampal | 0.143 | 0.0185 | 0.021 | 0.204 | 17 | 2.61 |
| Amygdala | 0.139 | 0.0595 | -0.005 | 0.224 | 17 | 2.02 |
| pericalcarine | 0.139 | 0.0352 | 0.008 | 0.210 | 17 | 2.29 |
| superiorparietal | 0.137 | 0.0155 | 0.023 | 0.191 | 17 | 2.69 |
| transversetemporal | 0.134 | 0.0322 | 0.010 | 0.198 | 17 | 2.33 |
| precuneus | 0.120 | 0.0139 | 0.021 | 0.159 | 17 | 2.74 |
| lateraloccipital | 0.110 | 0.2258 | -0.054 | 0.215 | 17 | 1.26 |
| posteriorcingulate | 0.108 | 0.0556 | -0.002 | 0.159 | 17 | 2.06 |
| insula | 0.103 | 0.2074 | -0.044 | 0.189 | 17 | 1.31 |
| medialorbitofrontal | 0.097 | 0.3308 | -0.074 | 0.207 | 17 | 1.00 |
| parsorbitalis | 0.096 | 0.2240 | -0.044 | 0.177 | 17 | 1.26 |
| isthmuscingulate | 0.093 | 0.1534 | -0.026 | 0.151 | 17 | 1.49 |
| lateralorbitofrontal | 0.090 | 0.1199 | -0.017 | 0.137 | 17 | 1.64 |
| Accumbens area | 0.081 | 0.1946 | -0.028 | 0.130 | 17 | 1.35 |
| parsopercularis | 0.054 | 0.6424 | -0.083 | 0.131 | 17 | 0.47 |
| caudalmiddlefrontal | 0.046 | 0.7500 | -0.085 | 0.116 | 17 | 0.32 |
| paracentral | 0.040 | 0.7987 | -0.067 | 0.086 | 17 | 0.26 |
| postcentral | 0.039 | 0.8060 | -0.067 | 0.085 | 17 | 0.25 |
| inferiorparietal | 0.026 | 0.8807 | -0.056 | 0.048 | 17 | -0.15 |
| precentral | 0.025 | 0.9125 | -0.105 | 0.095 | 17 | -0.11 |
| parstriangularis | 0.015 | 0.6906 | -0.096 | 0.065 | 17 | -0.40 |
| superiorfrontal | 0.012 | 0.7420 | -0.136 | 0.098 | 17 | -0.33 |
| supramarginal | 0.010 | 0.5336 | -0.088 | 0.047 | 17 | -0.64 |
| rostralmiddlefrontal | 0.009 | 0.5920 | -0.105 | 0.062 | 17 | -0.55 |

***Extended Data 5: Paired t-test results for regional alpha-predictive performance.*** *Model performances (correlation between predictions on held-out subjects and ground truth) when training on each bilateral region were compared to performance in the control condition which had the fMRI data shuffled in relationship to the EEG. Lower and upper limits refer to the confidence interval of the difference between region means and control.*

| Region | r | p-value | lower limit | upper limit | df | t-value |
|---|---|---|---|---|---|---|
| Putamen | 0.346 | 0.0001 | 0.060 | 0.155 | 31 | 4.57 |
| Caudate | 0.308 | 0.0011 | 0.030 | 0.108 | 31 | 3.60 |
| rostralanteriorcingulate | 0.307 | 0.0003 | 0.034 | 0.103 | 31 | 4.07 |
| caudalanteriorcingulate | 0.305 | 0.0023 | 0.026 | 0.108 | 31 | 3.33 |
| caudalmiddlefrontal | 0.292 | 0.0038 | 0.019 | 0.088 | 31 | 3.13 |
| posteriorcingulate | 0.289 | 0.0190 | 0.009 | 0.092 | 31 | 2.47 |
| superiorfrontal | 0.287 | 0.0077 | 0.014 | 0.083 | 31 | 2.85 |
| parsopercularis | 0.284 | 0.0096 | 0.012 | 0.079 | 31 | 2.76 |
| Thalamus | 0.276 | 0.0419 | 0.001 | 0.073 | 31 | 2.12 |
| insula | 0.273 | 0.0435 | 0.001 | 0.068 | 31 | 2.10 |
| fusiform | 0.269 | 0.0468 | 0.000 | 0.061 | 31 | 2.07 |
| Pallidum | 0.269 | 0.0733 | -0.003 | 0.063 | 31 | 1.85 |
| precentral | 0.268 | 0.0266 | 0.004 | 0.055 | 31 | 2.33 |
| lingual | 0.267 | 0.1031 | -0.006 | 0.062 | 31 | 1.68 |
| cuneus | 0.263 | 0.1252 | -0.007 | 0.057 | 31 | 1.58 |
| supramarginal | 0.263 | 0.1042 | -0.005 | 0.053 | 31 | 1.67 |
| transversetemporal | 0.263 | 0.1169 | -0.006 | 0.055 | 31 | 1.61 |
| Amygdala | 0.262 | 0.0969 | -0.004 | 0.051 | 31 | 1.71 |
| rostralmiddlefrontal | 0.260 | 0.2816 | -0.019 | 0.062 | 31 | 1.10 |
| Accumbens area | 0.259 | 0.1414 | -0.007 | 0.049 | 31 | 1.51 |
| pericalcarine | 0.258 | 0.1263 | -0.006 | 0.044 | 31 | 1.57 |
| middletemporal | 0.250 | 0.2264 | -0.008 | 0.031 | 31 | 1.23 |
| parstriangularis | 0.250 | 0.3378 | -0.012 | 0.034 | 31 | 0.97 |
| superiorparietal | 0.249 | 0.3831 | -0.014 | 0.035 | 31 | 0.88 |
| inferiortemporal | 0.249 | 0.3035 | -0.010 | 0.030 | 31 | 1.05 |
| parsorbitalis | 0.248 | 0.5397 | -0.021 | 0.039 | 31 | 0.62 |
| precuneus | 0.247 | 0.3962 | -0.011 | 0.027 | 31 | 0.86 |
| lateraloccipital | 0.245 | 0.5907 | -0.016 | 0.028 | 31 | 0.54 |
| superiortemporal | 0.242 | 0.6368 | -0.011 | 0.018 | 31 | 0.48 |
| postcentral | 0.241 | 0.6761 | -0.011 | 0.016 | 31 | 0.42 |
| bankssts | 0.241 | 0.7664 | -0.016 | 0.021 | 31 | 0.30 |
| parahippocampal | 0.241 | 0.7862 | -0.016 | 0.021 | 31 | 0.27 |
| Hippocampus | 0.240 | 0.8425 | -0.016 | 0.019 | 31 | 0.20 |
| inferiorparietal | 0.239 | 0.9505 | -0.023 | 0.024 | 31 | 0.06 |
| isthmuscingulate | 0.239 | 0.9260 | -0.013 | 0.014 | 31 | 0.09 |
| lateralorbitofrontal | 0.236 | 0.7668 | -0.022 | 0.017 | 31 | -0.30 |
| paracentral | 0.231 | 0.2418 | -0.021 | 0.006 | 31 | -1.19 |
| medialorbitofrontal | 0.227 | 0.4080 | -0.039 | 0.016 | 31 | -0.84 |

***Extended Data 6: Paired t-test results for regional delta-predictive performance.*** *Model performances (correlation between predictions on held-out subjects and ground truth) when training on each bilateral region combined with non-gray matter regions were compared to performance in the control condition which had only the non-gray matter regions. Lower and upper limits refer to the confidence interval of the difference between region means and control.*

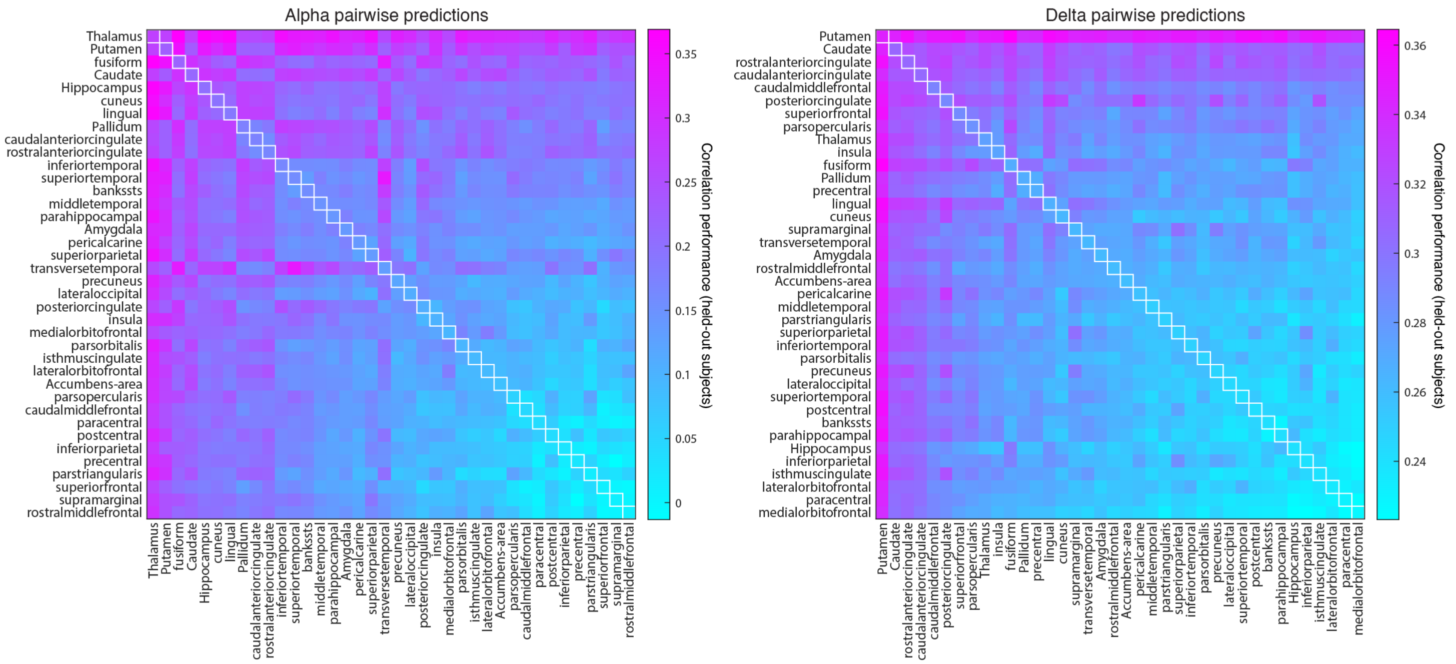


***Extended Data 7: Distinct prediction patterns when model is trained on every possible combination of bilateral gray matter regions.*** *For delta predictions, the non-gray matter regions were also included. Regions are ordered based on performance in the individual regions analysis (Fig. 3 and Extended Data 4-6).*

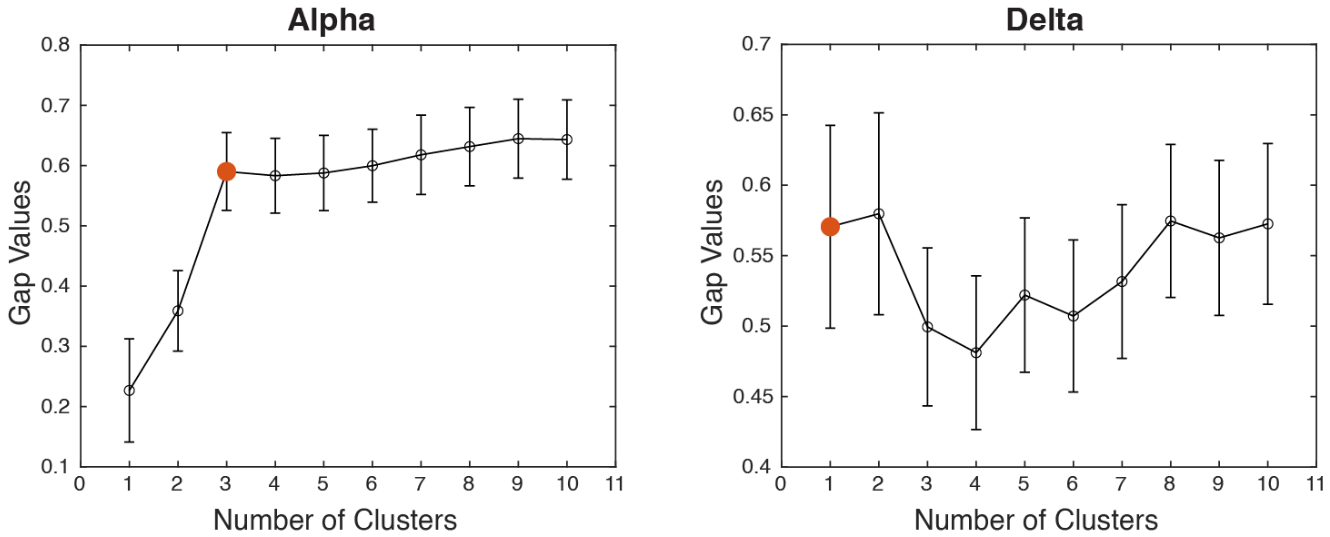


***Extended Data 8: Clustering analysis reveals the optimal number of clusters that explain pairwise performance benefits for each frequency band*** *(red circle). The gap statistic was used to evaluate each clustering solution for alpha- and delta-predictive information. See methods for details on how the optimal gap value is determined.*

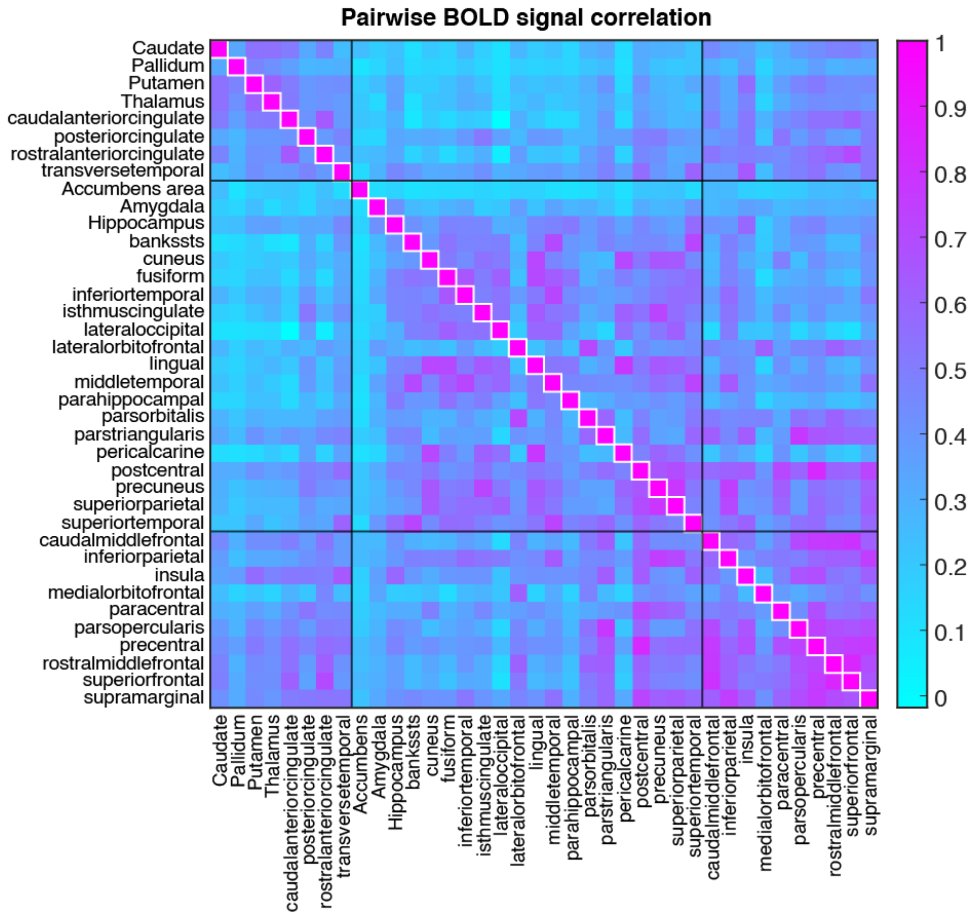


***Extended Data 9: Decoded information results in highly discrete networks (see Fig. 4), and only weakly reflects correlation between the BOLD activity of different brain regions.*** *A Pearson's correlation value between the BOLD signals of each pair of brain regions (similar as what is done in resting state functional connectivity analyses) was calculated for each subject used in the alpha clustering analysis. To determine that clustering behavior is a result of shared decoded information, and not only BOLD signal correlation, the mean pairwise BOLD correlation values were plotted following the clustering structure shown in Fig. 4b. Compared to the highly separable results shown in Fig. 4, the pairwise correlation values of the BOLD signal only weakly reflects the identified clusters.*

*.*

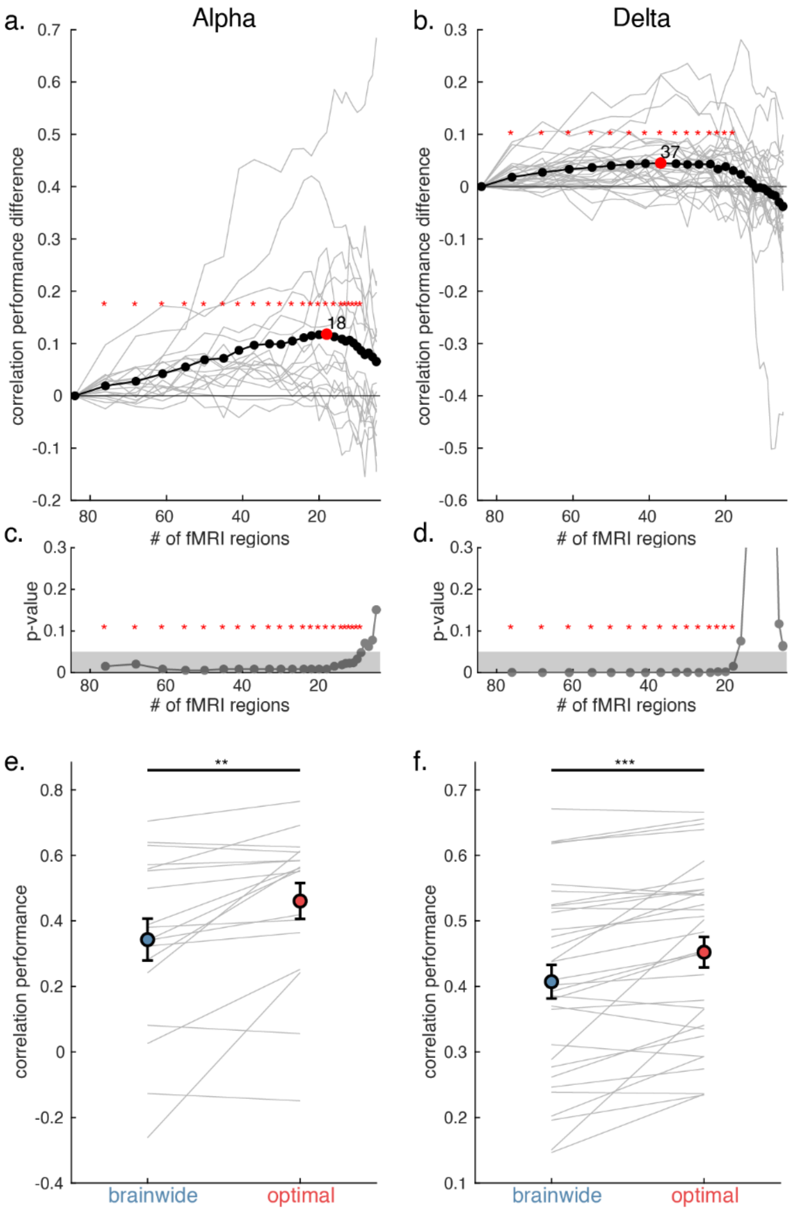


***Extended Data 10: Alpha rhythms can be predicted from a small spatial network, whereas delta requires information from a broader set of regions.*** *Iteratively removing fMRI regions associated with low weights increased model performance up until a plateau, with different spatial scales associated with the best alpha and delta predictions.* ***a.*** *Alpha prediction performance peaked at 18 regions. The lowest-weight fMRI regions at each iteration were removed from the model, and then training and testing were performed again for the next iteration. Gray lines represent individual subjects. The iteration with best performance is marked in red. Red stars indicate iterations with significant improvements (as compared to all regions). Multiple comparisons controlled using the Benjamini-Hochberg procedure.* ***b.*** *Delta prediction performance, equivalent to panel a, showing performance peaking at 37 regions.* ***c.*** *P-values (from two-tailed t-tests, corrected using the Benjamini-Hochberg procedure) of alpha performance improvements relative to using all regions, showing that improvements cease to be significant when model contains 8 or fewer fMRI regions.* ***d.*** *P-values of delta performance improvements, equivalent to panel c, showing that improvements cease to be significant when model contains 17 or fewer regions.* ***e.*** *Mean alpha prediction performance when using brainwide information vs. when using the optimal number of regions identified in panel a demonstrates significant improvements with feature selection. ** p<0.01. Error bars are SEM; gray lines are individual subjects.* ***h.*** *Delta prediction performance also demonstrates significant improvements with feature selection. *** p<0.001.*

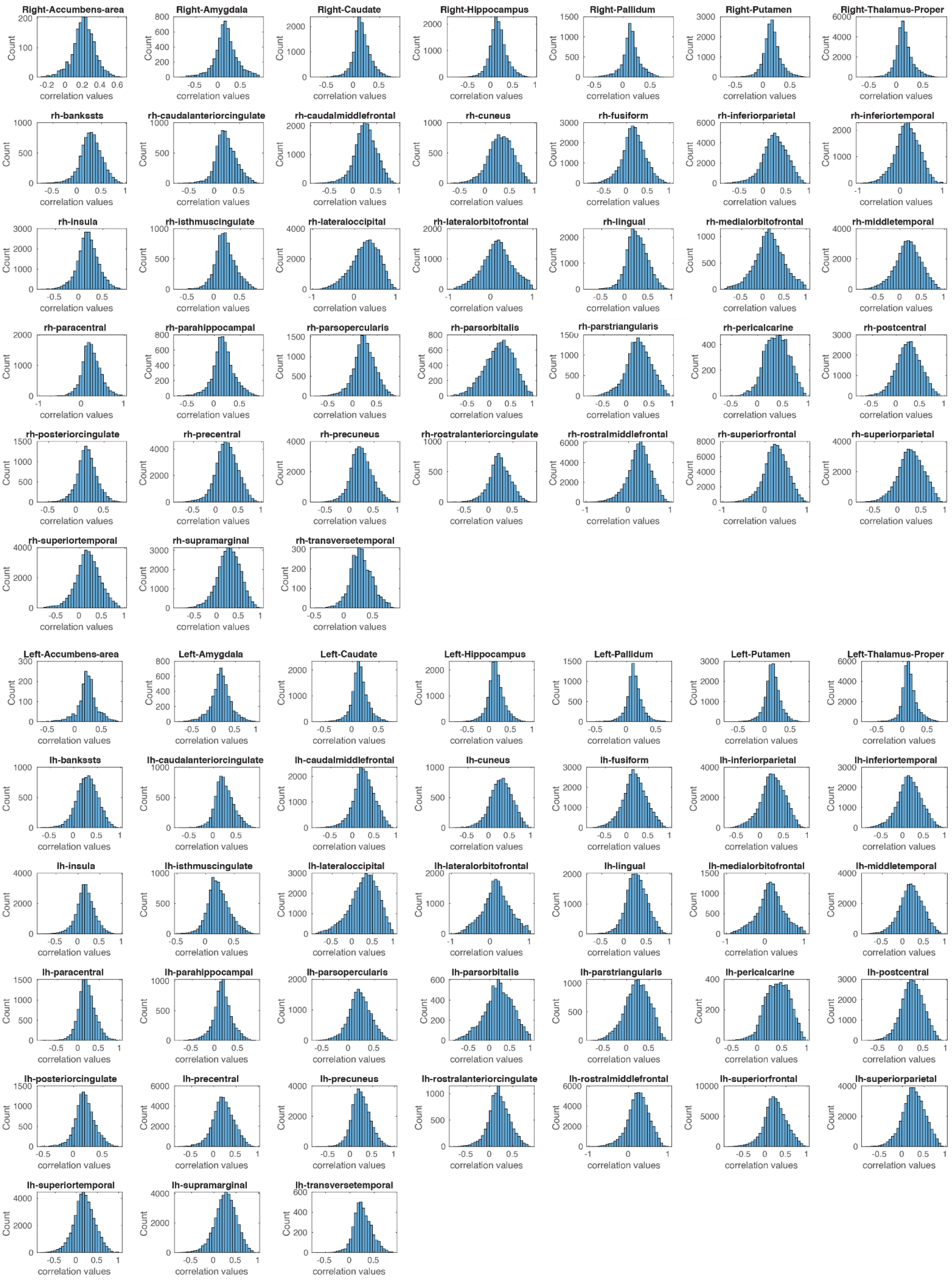
Right-Accumbens-area
Right-Amygdala
Right-Caudate
Right-Hippocampus
Right-Pallidum
Right-Putamen
Right-Thalamus-Proper
rh-bankssts
rh-caudalanteriorcingulate
rh-caudalmiddlefrontal
rh-cuneus
rh-fusiform
rh-inferiorparietal
rh-inferiortemporal
rh-insula
rh-isthmuscingulate
rh-lateraloccipital
rh-lateralorbitofrontal
rh-lingual
rh-medialorbitofrontal
rh-middletemporal
rh-paracentral
rh-parahippocampal
rh-parsopercularis
rh-parsorbitalis
rh-parstriangularis
rh-pericalcarine
rh-postcentral
rh-posteriorcingulate
rh-precentral
rh-precuneus
rh-rostralanteriorcingulate
rh-rostralmiddlefrontal
rh-superiorfrontal
rh-superiorparietal
rh-superiortemporal
rh-supramarginal
rh-transversetemporal
Left-Accumbens-area
Left-Amygdala
Left-Caudate
Left-Hippocampus
Left-Pallidum
Left-Putamen
Left-Thalamus-Proper
lh-bankssts
lh-caudalanteriorcingulate
lh-caudalmiddlefrontal
lh-cuneus
lh-fusiform
lh-inferiorparietal
lh-inferiortemporal
lh-insula
lh-isthmuscingulate
lh-lateraloccipital
lh-lateralorbitofrontal
lh-lingual
lh-medialorbitofrontal
lh-middletemporal
lh-paracentral
lh-parahippocampal
lh-parsopercularis
lh-parsorbitalis
lh-parstriangularis
lh-pericalcarine
lh-postcentral
lh-posteriorcingulate
lh-precentral
lh-precuneus
lh-rostralanteriorcingulate
lh-rostralmiddlefrontal
lh-superiorfrontal
lh-superiorparietal
lh-superiortemporal
lh-supramarginal
lh-transversetemporal
Count
correlation values

***Extended Data 11: Correlation between individual voxels within a region, and the regional mean timeseries, across all subjects and runs.*** *In each subject and run, the voxels within each anatomical region were averaged to generate the region timeseries. This averaging process resulted in a mean timeseries that was well-correlated with most voxels in a region, providing dimensionality reduction and signal-to-noise ratio improvement, in addition to being a way to bridge across different subjects' anatomies. However, some voxels in each regions were not well captured by the mean. These voxels could contain additional information that would require a more complex model to uncover.*
